# Differential functional coupling in Gp130-JAK complexes expands the plasticity of the interleukin-6 signaling axis

**DOI:** 10.1101/2023.05.24.542077

**Authors:** Alison McFarlane, Junel Sotolongo Bellón, Thomas Meyer, Elizabeth Pohler, Jacob Piehler, Ignacio Moraga

## Abstract

Cytokines dimerize/oligomerize surface receptors to activate signaling. While cytokine receptors preferentially bind only one member of the JAK family, ancestral cytokine receptors, such as Gp130, promiscuously recruit different JAKs to elicit their activities. Here, we have explored how the identity of JAKs in Gp130 signaling complexes can regulate functional outcomes. Using a synthetic biology approach, we show that Gp130 bound to different JAKs propagates distinct STAT activation profiles. While Gp130-JAK1 complexes activated both, STAT1 and STAT3 very potently, Gp130-JAK2 complexes exhibited a clear preference for STAT3 activation. Gp130-TYK2 complexes triggered overall weaker signaling but with diminished STAT specificity. The three JAKs competed for binding to Gp130 and led to differential activation of phospho-Tyr in the Gp130 intracellular domain. JAK1, JAK2 and to a lower extent TYK2 bound with comparable affinities to Gp130, and in response to IL-6 stimulation efficiently drove Gp130 dimerization. However, the three JAKs differentially affected Gp130 surface expression, identifying JAK-dependent receptor trafficking as a critical determinant of signaling plasticity. Our results provide new mechanistic insights into how differential functional coupling in Gp130-JAK complexes translates into unique signaling signatures that likely contribute to its large functional diversity.

## INTRODUCTION

Cytokines control all aspects of mammalian physiology (*1–3*). Deregulation of cytokines or cytokine-induced signaling networks often results in disease, making this family highly relevant for human health (*4, 5*). Cytokines of the class I/II family dimerize/oligomerize cell surface receptors to trigger the activation of the JAK (Janus kinase)/STAT (signal transducer and activator of transcription) signaling pathway but can also control the activity of diverse serine/threonine kinase signaling networks (*6, 7*). The canonical view of cytokine signaling describes a linear process where upon formation of the cytokine-receptor complex, JAKs are activated via transphosphorylation and drive the phosphorylation of Tyr residues in the cytokine receptor intracellular domain (*7–10*). Phosphorylated Tyr residues (pTyr) then act as docking sites for STATs, which upon binding, are phosphorylated by JAKs, and form homo and/or hetero-dimers that translocate to the nucleus and induce specific gene programs (*11, 12*).

With four different JAKs (JAK1, JAK2, JYK3 and TYK2) and seven STATs (STAT1-STAT4, STAT5a/b and STAT6), diverse combinations of JAKs and STATs are found among different cytokine receptors. In most cases, specific JAKs are assigned to each cytokine receptor subunit, which typically promote the activation of a set of defined STAT members to exert unique responses (*11*). In recent years, however, it has become apparent that cytokine signaling is a more dynamic and plastic process that allows cells to adapt their responses to the changing environment (*6*). STATs compete for binding to shared pTyr motifs in the cytokine receptor intracellular domains (*13*). This competition makes cytokine signaling signatures very sensitive to changes in STAT concentration. For instance, IL-10 loses its anti-inflammatory properties in cells with high STAT1 expression levels, because of a pro-inflammatory environment rich in IFNs (*14*). Increased STAT1 levels outcompetes STAT3 for binding to the different cytokine receptors, despite binding these receptors with lower affinity. This STAT competition is critical in defining cytokine responses in disease. We recently showed that STAT1 expression levels are increased in Crohn’s disease and SLE patients and contribute to alter IL-6 activities (*15*). Interestingly, this dynamic equilibrium between signaling molecules and cytokine receptors is not limited to STATs. Some cytokine receptors can bind more than one type of JAK in a competitive manner (*16–18*). How the identity of the receptor-JAK complex influence cytokine responses, however, remains less well understood.

The different JAKs/STATs components engaged by cytokines to elicit their activities are well characterized (*7–10*). How JAK/STAT proteins integrate into a signaling network capable of producing high pleiotropy yet specific functional responses however requires further investigation. The wide expression of the JAK/STAT proteins across most cell types in conjunction with the presence of diverse regulators of JAK/STAT signaling has hindered the study of the contribution of each component of the network to a given bioactivity. Synthetic biology and pathway rebuilding approaches provide a unique opportunity to study how the different components of the cytokine signaling network coordinate to produce specific responses (*19*). Rebuilding the JAK/STAT pathway one component at a time could help uncovering how each component of this pathway contributes to the overall functional outcome. This approach has been used in the past for other ligand-receptor systems, such as the B cell receptor (*20*), where it has provided critical molecular insights into the functioning and biology of these systems.

IL-6 is an ideal model system to address how signaling specificity is achieved by cytokine receptors. IL-6 exhibits a high degree of pleiotropic activities ranging from pro- to anti-inflammatory responses and acts as a central regulator of the immune response (*21–24*). IL-6 activates signaling by engaging two surface receptors, IL-6Rα and Gp130, and forming a hexameric complex on the surface of responsive cells (*25*). This IL-6 hexameric complex triggers the activation of the JAK/STAT1/STAT3 signaling pathway (*26*). While JAK1 has been described to be the dominant JAK associated with Gp130, JAK2 and TYK2 can also bind this receptor (*16, 17*). How the identity of the Gp130-JAK complex influences functional outcome remains poorly understood. Here, we systematically explored how differential JAK usage expands the plasticity of Gp130 signaling. To this end, we reconstituted IL-6-Gp130 signaling pathway in *Drosophila* Schneider Cells (S2 cells) to gain molecular insights into how Gp130 translates extracellular cytokine binding information into specific signaling responses, unbiased from the many additional regulatory mechanism present in mammalian cells. Importantly, human cytokine receptors, JAKs and STATs are not activated by the only related endogenous *Drosophila* receptor dome, allowing the rebuilding of the Gp130 JAK/STAT pathway from scratch. We have systematically explored how Gp130 integrates signaling responses when bound to independent JAK and STAT members. Quantitative signaling studies revealed that the identity of the Gp130-JAK complex critically contributes to define signaling signatures. While Gp130-JAK1 complexes activate STAT1/STAT3 very potently, with other STATs being activated less efficiently, Gp130-JAK2 complexes exhibit a clear preference for STAT3 activation, leading to higher levels of STAT3 phosphorylation than those induced by Gp130-JAK1 complexes. Gp130-TYK2 complexes overall activate weaker signaling with a preference for STAT3 activation as well. JAK1, JAK2 and TYK2 compete for binding to Gp130, and lead to differential activation of phospho-Tyr residues in the Gp130 intracellular domain. Interestingly, in HeLa cells Gp130-JAK2 and Gp130-TYK2 complexes triggered signaling less efficiently than Gp130-JAK1 complexes. These differences could not be explained by reduced expression levels of JAK2 and TYK2 in HeLa cells or by their binding affinity to Gp130, suggesting that alternative mechanisms are in place in HeLa cells to limit Gp130-JAK2/TYK2 signaling. While effective, ligand-induced Gp130 dimerization was confirmed for JAK1, JAK2 and TYK2, we found that JAKs differentially affect Gp130 cell surface expression, pointing towards altered endocytosis and/or endocytic trafficking. Our results provide new molecular insights into how JAKs differentially regulate Gp130 signaling at multiple levels to elicit unique signaling signatures that likely contribute to its large functional diversity. At the more practical level, our results suggest that by altering their JAK expression levels, cells could radically change their responses to any given cytokine, leading to high degree of functional plasticity.

## RESULTS

### The Gp130 signaling pathway can be reconstituted in S2 Cells

How cytokine signaling networks integrate cytokine-receptor binding into highly pleiotropic responses remains poorly understood. To address the role of differential JAK usage, we have used *Drosophila* S2 cells to reconstruct the human Gp130 JAK/STAT pathway. Initially, we assessed whether S2 cells would be a suitable system to detect IL-6 mediated signaling responses. S2 cells were transfected with Gp130, JAK1 and STAT3 via nucleofection (Fig. 1A). To monitor transfection efficiency, we tagged Gp130 with mEGFP in its N-term. Transfection efficiencies between 50-70% were obtained routinely (Fig. 1B). Importantly, Gp130, JAK1 and STAT3 could be detected simultaneously in S2 cells, indicating the ability of these cells to express all the components required to trigger signaling in response to IL-6 stimulation (Fig. 1B). Previous studies have shown that JAK overexpression can lead to STAT phosphorylation in a ligand-independent manner (*27*). Thus, to limit ligand-independent JAK/STAT activation in S2 cells, which could confound our observations, we performed a JAK1 titration experiment to find the concentration of JAK1 that would result in maximal ligand-induced STAT3 activation without basal activity. 5 μg of GFP-Gp130 and STAT3 plasmids were transfected together with different concentrations of JAK1 DNA. Transfection of more than 3 μg of JAK1 cDNA produced a significant increase in the ligand-independent pSTAT3 phosphorylation levels (Fig. S1A). Based on this, we decided to use 3 μg of JAK1 for the subsequent experiments.

**Figure 1.**
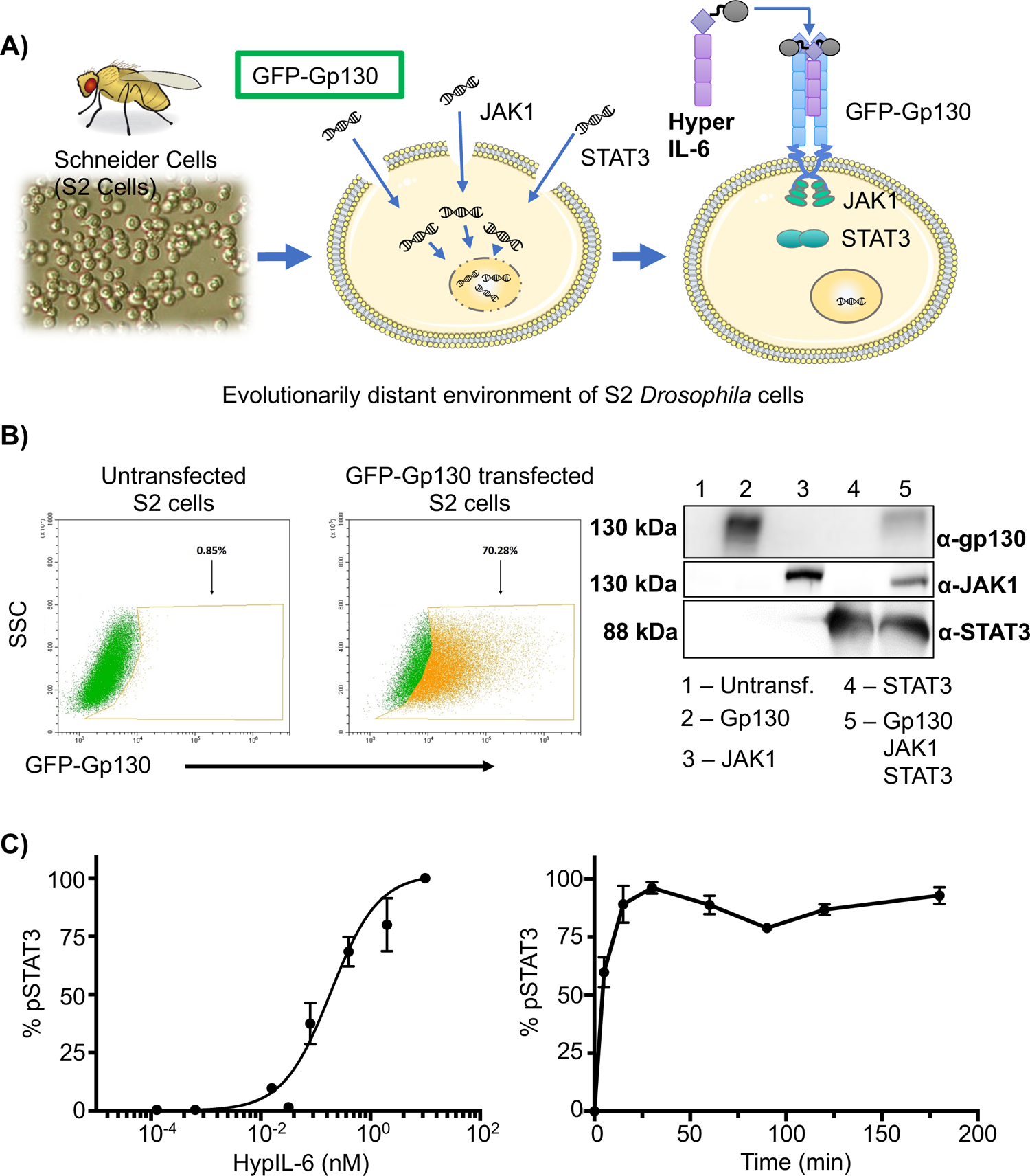
Drosophila S2 cells can express all components of the IL-6-Gp130 signaling pathway. **A)** Workflow scheme of the S2 cell transfection system. **B)** S2 cells were nucleofected with the different components of the IL-6-Gp130 signaling pathway and their expression was monitored by flow cytometry and western blotting. Left panel shows GFP-Gp130 transfected S2 cells with > 50% transfection efficiency. Right panel shows western blot of S2 cells nucleofected with GFP-Gp130/JAK1/STAT3. S2 cells were able to successfully incorporate all components of the Gp130/JAK/STAT pathway. An untransfected control was also included. **C)** S2 cells were nucleofected with GFP-Gp130, JAK1 and STAT3 and then stimulated with a 1:5 serial dilution of HypIL-6 with a starting concentration of 20 nM (left panel); or with 20 nM of HypIL-6 for 0-180 minutes (right panel). Data are mean +/- SEM from three independent replicates.

To assess the characteristics of STAT3 phosphorylation induced in response to IL-6 stimulation in transfected S2 cells, we performed dose-response and kinetics studies. We have used Hyper-IL-6 (HypIL-6) for stimulation (*28*), since this form of IL-6 does not require expression of the IL-6Rα receptor for signaling, thus simplifying our transfection panel. A 1:5 dilution series was prepared of HypIL-6 stimulus. S2 cells transfected with GFP-Gp130/JAK1/STAT3 constructs were incubated with the varying concentrations of HypIL-6 for 15 minutes and STAT3 phosphorylation levels were assayed by phospho-flow cytometry. HypIL-6 stimulation results in a concentration-dependent activation of STAT3-Tyr705 (Fig. 1C). Kinetic time course studies showed that pSTAT3-Tyr705 levels induced by HypIL-6 in S2 cells increased very rapidly, peaking at 15 min, and remained at a maximal level throughout the time course examined (Fig. 1C, right panel and Fig. S1B). This is in stark contrast to previous observations in mammalian cells. Here, pSTAT3-Tyr705 by IL-6 peaks at 15 min and gradually decreases to basal levels after 3-5 hours (*15, 29*). Inhibition of JAK1 activity using the JAK inhibitor tofacitinib (TOFA) resulted in an acute decrease in pSTAT3-Tyr705 levels (Fig. S1C), indicating that JAK activity is required for sustained STAT3 phosphorylation in S2 cells and suggesting that endogenous S2 cells phosphatases can dephosphorylate human STAT3. These observations thus suggest that lack of negative feedback mechanisms in S2 cells result in sustained STAT3 phosphorylation. Based on these observations, we decided to focus on dose-response studies at 15 min stimulation for the rest of this study. Overall, these data validate that the human JAK/STAT components are functioning within S2 cells which verified the suitability of this system for the molecular analysis of the human Gp130 JAK/STAT pathway.

### The identity of the Gp130-JAK complex defines STAT activation by IL-6

Gp130 has been reported to engage JAK1, JAK2 and TYK2 to activate signaling (*16, 17*). However, whether the identity of the Gp130-JAK complex defines unique phospho-STAT (pSTAT) activation profiles that contribute to Gp130 functional heterogeneity is currently not known. To tackle this question, we investigated STAT activation profiles induced by different Gp130-JAK complexes. For this purpose, we transfected S2 cells with GFP-Gp130 and then combined each JAK (JAK1/JAK2/JAK3/TYK2) with each STAT (STAT1-6) to generate a 6×4 matrix of possible JAK/STAT combinations (Fig. 2A). A series of detailed dose-response experiments using HypIL-6 were performed to assess pSTAT activation profiles induced by Gp130-JAK complexes (Fig. 2B and Fig. S2). As expected, Gp130 triggered STAT phosphorylation when bound to JAK1, JAK2 or TYK2 but not when bound to JAK3, confirming the specificity of this latter for the common gamma chain (Fig. 2B and Fig. S2). Gp130-JAK1 complexes triggered potent pSTAT1 and pSTAT3 activation, with pSTAT4 and pSTAT5 proteins being activated to a lower extent (Fig. 2B and 2C and Fig. S2). pSTAT2 and pSTAT6 were not activated by this complex. Importantly, all JAKs and STATs were similarly well expressed in S2 cells (Fig. S3). Interestingly, Gp130-JAK2 complexes also led to potent pSTAT activation, but the identity of pSTATs yielded by this complex differed significantly from those induced by Gp130-JAK1 complexes. Gp130-JAK2 complexes phosphorylated STAT3 more potently than Gp130-JAK1 complexes, but showed much weaker activation of the other STAT proteins (Fig. 2B and 2C and Fig. S2). Gp130-TYK2 complexes activated pSTAT profiles similar to those induced by Gp130-JAK2 complexes, with slightly better activation of pSTAT4 (Fig. 2B and 2C and Fig. S2). Overall, these results suggest that the identity of the Gp130-JAK complex instructs specific pSTAT patterns that could impact Gp130 biological responses. Since JAK- and STAT-specific regulatory feedback mechanisms can be largely ruled out by this experimental approach, we conclude that some selectivity of STAT phosphorylation is encoded by different JAKs and their structural organization in Gp130 dimers.

**Figure 2.**
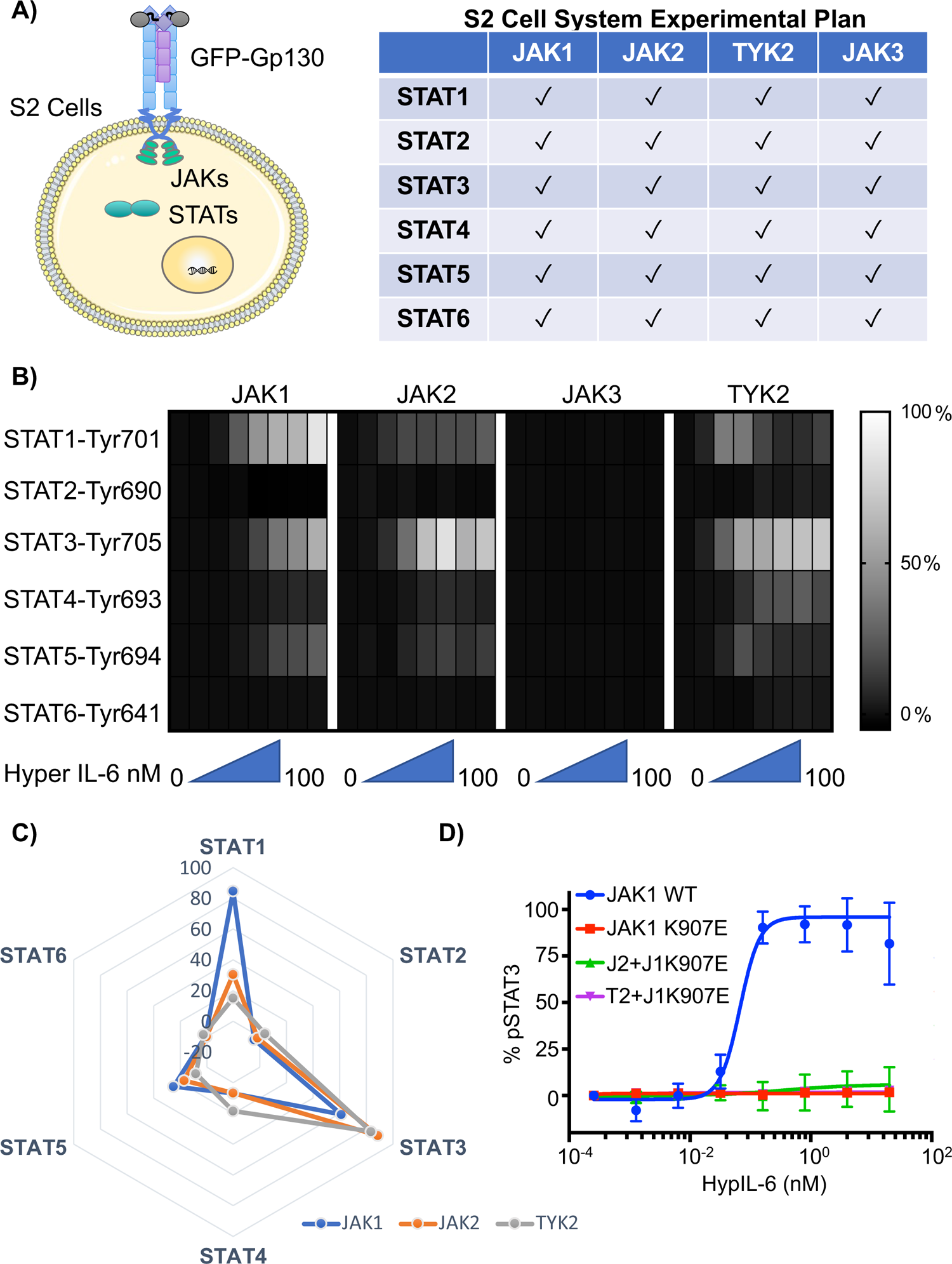
JAK1-, JAK2- and TYK2-Gp130 complexes trigger different profiles of STAT activation in S2 cells. **A)** Schematic representation and table of the combinatorial matrix designed to explore all signaling combinations compatible with Gp130 engagement. **B)** Heatmap of resulting dose response experiments following 15 min stimulation with HypIL-6 showing the percentage of pSTAT activation. Each box in this heatmap represents the normalized mean fluorescence intensity values (normalized to JAK1). Data are mean +/- SEM from three independent experiments. **C)** Filled radar representation of the signaling molecules activated by the different Gp130-JAKs complexes. The STAT molecules activated by the ligands are shown on the perimeter of the circle and their respective activation potencies are denoted by the radius of the circle. The different shapes of the filled radar exhibited by the different ligands define their distinct signaling signatures. **D)** Dose response experiments after stimulation with HypIL-6 stimulation in S2 cells transfected with JAK1 WT, the dominant negative JAK1 K907E mutant, or the JAK1 K907E mutant along with JAK2 or TYK2. JAK1 K907E abolished the pSTAT3 signal of both JAK2 and TYK2, proving JAKs bind competitively to Gp130. Data are mean +/- SEM from three independent experiments.

The molecular details defining Gp130-JAK interaction are still under debate. While early studies showed that in the absence of JAK1, the phosphorylation of STAT1/STAT3 upon activation with IL-6 is greatly reduced, a kinase-negative mutant of JAK2 inhibited IL-6 mediated signaling activation, suggesting that JAK1 and JAK2 compete for binding to Gp130 (*30*). To address this, we generated a dominant negative JAK1 mutant - JAK1-K907E - containing a BFP-tag on its C-terminus, allowing easy tracking via cytometric analysis. The K907E mutation is located within the kinase domain of JAK1 and has been shown to abolish JAK1 catalytic activity (*31, 32*). However, the mutation does not interfere with JAK1’s ability to bind to the cytokine receptor (*31, 32*). Co-expression of the JAK1-K907E mutant with other JAKs, prevented HypIL-6 mediated signaling in the JAK1, JAK2 and TYK2 backgrounds, confirming that JAKs are competing for binding on the Gp130 receptor (Fig. 2D).

### Gp130-JAKs complexes exhibit different requirements for Tyr usage in Gp130

The identity of the Gp130-JAK complex influences the profile of STAT molecules activated. We hypothesize that different Gp130/JAK complexes engage unique Tyr pools in the Gp130 intracellular domain (ICD) resulting in differential STAT activation. Gp130 has six Tyr residues in its ICD that can be potentially phosphorylated by JAKs and thus promote STAT recruitment. We generated a series of Gp130 mutants where ICD Tyr were mutated to Phe to prevent their phosphorylation (Fig. 3A). In-depth dose-response studies were performed to assess signaling potential by Gp130mut-JAK complexes. S2 cells were transfected with GFP-Gp130 or GFP-Gp130 mutants in combinations with JAK1, JAK2 or TYK2 and STAT1 or STAT3. Transfected cells were then stimulated with the indicated doses of HypIL-6 and analyzed via phospho-flow cytometry for pSTAT1-Tyr701 or pSTAT3-Tyr705 levels (Fig. 3B and Fig. S4A). Gp130-JAK1 complexes triggered robust STAT3 phosphorylation even when three out of the six Gp130 ICD Tyr were mutated to Phe (Fig. 3B and Fig. S4A). Mutation of more than three Tyr led to a dramatic decrease in STAT3 phosphorylation levels, which dropped to 20% of levels induced by wt Gp130 (Fig. 3B and Fig. S4A). Interestingly, even in the absence of ICD Tyr, Gp130 could still trigger low but detectable STAT3 phosphorylation upon HypIL-6 stimulation (Fig. S4A). STAT1 phosphorylation was more sensitive to changes in the number of Gp130 ICD Tyr supporting our previous observations (*29*). Mutation of three Tyr on Gp130 resulted in a drop of 50% in STAT1 phosphorylation levels induced by HypIL-6 (Fig. 3B and Fig. S4A). Mutation of more than three Tyr on Gp130 resulted in very weak STAT1 phosphorylation by HypIL-6. As for STAT3, Gp130 could trigger very low but detectable levels of STAT1 phosphorylation even in the complete absence of phosphorylated Tyr (Fig. 3B and Fig. S4A).

**Figure 3.**
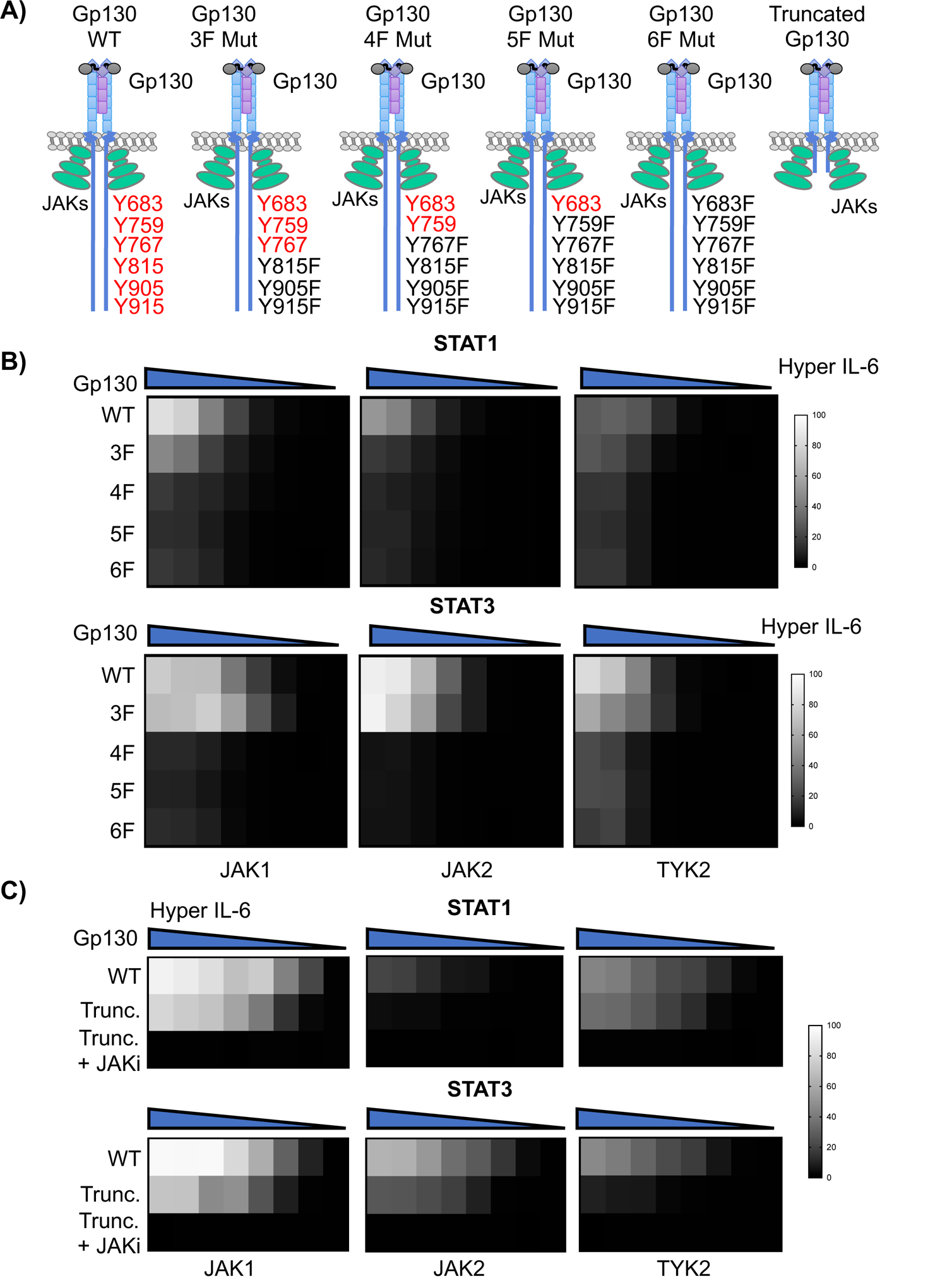
Different usage of Gp130 intracellular Tyr by the Gp130-JAK complexes. A) Schematic representation of Gp130 mutants generated showing tyrosines targeted for inactivation. B) Heatmap of resulting dose response experiments following 15 min stimulation with HypIL-6 showing the percentage of pSTAT1/pSTAT3 activation for Phe-Gp130 mutants. Each box in this heatmap represents the normalized mean fluorescence intensity values (normalized to JAK1). Data are mean + /- SEM from three independent experiments. **C)** Heatmap of resulting dose response experiments following 15 min stimulation with HypIL-6 showing the percentage of pSTAT1/pSTAT3 activation for the Gp130 truncated mutant. The JAK inhibitor (JAKi) Tofacinitib was used to assess JAK activity contribution to STAT phosphorylation. Each box in the heatmap represents the normalized mean fluorescence intensity values (normalized to JAK1). Data are mean + /- SEM from three independent experiments.

For Gp130-JAK2 and Gp130-TYK2 complexes, we observed noticeable differences in their signaling capabilities, as compared to Gp130-JAK1 complexes, upon mutation of the tyrosine sites on Gp130. As Gp130-JAK1 complexes, Gp130-JAK2 complexes triggered potent STAT3 phosphorylation when three Tyr were mutated in Gp130 (Fig. 3B and Fig. S4A). Mutation of more than three Tyr resulted in significant loss of STAT3 phosphorylation. However, the mutation of three Tyr resulted in loss of STAT1 phosphorylation by Gp130-JAK2 complexes (Fig. 3B and Fig. S4A). This led to a strong pSTAT3 bias by Gp130-JAK2 complexes when Gp130 ICD Tyr were limited (Fig. S4B). Contrary to Gp130-JAK1 and Gp130-JAK2 complexes, Gp130-TYK2 complexes could sense the loss of three Tyr in the Gp130 ICD, with levels of STAT3 phosphorylation decreasing by 20% in this context (Fig. 3B and Fig. S4A). Interestingly, even though Gp130-TYK2 complexes triggered very weak STAT1 phosphorylation, these activation levels were minimally affected when Gp130 ICD Tyr were mutated to Phe (Fig. 3B and Fig. S4A). Overall, our results suggest that different Gp130-JAK complexes access different pools of Gp130 ICD Tyr to activate unique STATs profiles.

### Truncated Gp130 mutants retain JAK-dependent pSTAT activation patterns

We were surprised to find that Gp130 mutants lacking tyrosine residues in their intracellular domains still promoted low but significant STAT1/STAT3 phosphorylation. This residual pSTAT activation could result from STATs binding the Gp130 ICD even in the absence of phosphorylated Tyr, or from STATs using JAKs as scaffolds for their binding and activation. To distinguish between these two possibilities, we designed a new Gp130 mutant where the intracellular domain was truncated (Fig. 3A). The truncated Gp130 mutant was carefully designed to ensure that the Box 1 (‘WPNVPDPS’) and Box 2 (‘DVSVVEIEAN’) motifs, which are essential for JAK binding, remained intact. S2 cells were transfected with wt GFP-Gp130 or the truncated GFP-Gp130 mutant, along with the indicated JAK and STATs, stimulated with HypIL-6 and probed for pSTAT1-Tyr701 and pSTAT3-Tyr705 via phospho-flow cytometry (Fig. 3C and Fig. S4D). We included the JAK inhibitor TOFA as a negative control to ascertain the contribution of JAKs in STAT phosphorylation. When we analyzed expression of the different Gp130 variants, we noticed that the truncated Gp130 mutant expressed significantly better on the surface of the S2 cells than WT Gp130 (Fig. S4C). Surprisingly, signaling by the truncated Gp130 construct closely resembled that obtained by Gp130 wt, only showing a small decrease in signaling potency (Fig. 3C and Fig. S4D). Importantly, inhibition of JAK activity, via TOFA treatment, resulted in complete loss of STAT1/STAT3 phosphorylation, highlighting that JAKs are responsible for STAT activation by the truncated Gp130 mutant (Fig. 3C and Fig. S4D). These data suggest that STAT may interact directly with JAKs, thus bypassing the need of receptor intracellular Tyr for their activation. Interestingly, we still observed similar, JAK-dependent pSTAT1/pSTAT3 patterns as for wt Gp130, corroborating substrate specificity is encoded by the identity of Gp130-JAK signaling complexes.

### GP130 binds JAK1 and JAK2 with similar affinity

Altered Gp130 signaling by JAK2 and TYK2 as compared to JAK1 could be caused by differences in efficiency and dynamics of JAK recruitment to the receptor. To test this hypothesis, we probed JAK interaction with Gp130 in the plasma membrane by live cell micropatterning (*33, 34*). To this end, HeLa cells expressing Gp130 N-terminally fused to a HaloTag and mTagBFP (HaloTag-BFP-Gp130) were cultured on substrates presenting micropatterned HaloTag ligand (HTL, Fig. 4A). This led to the capture and spatially reorganization of HaloTag-BFP-Gp130 in the plasma membrane of live cells. To assess binding efficiencies between different JAKs and Gp130, we co-expressed the FERM/SH2 domain of JAK1, JAK2 or TYK2 fused to mEGFP (JAK-FS). The JAK-FS domain is responsible for JAK recruitment by interacting with box 1 and box 2 motifs of the receptor and therefore is ideal to selectively probe JAK-Gp130 interaction (*35*). JAKs interacting with Gp130 would be expected enrich the Gp130 micropatterns depending on their binding affinity (Fig. 4A). The relative binding affinity was estimated from the relative contrast of the micropatterns observed in the GFP channel upon imaging by total internal reflection fluorescence (TIRF) microscopy (Fig. 4B, C). While the contrast was very similar for JAK1 and JAK2, TYK2 showed significantly less binding. For JAK3, binding was just above the negative control, suggesting very weak Gp130 binding.

**Figure 4.**
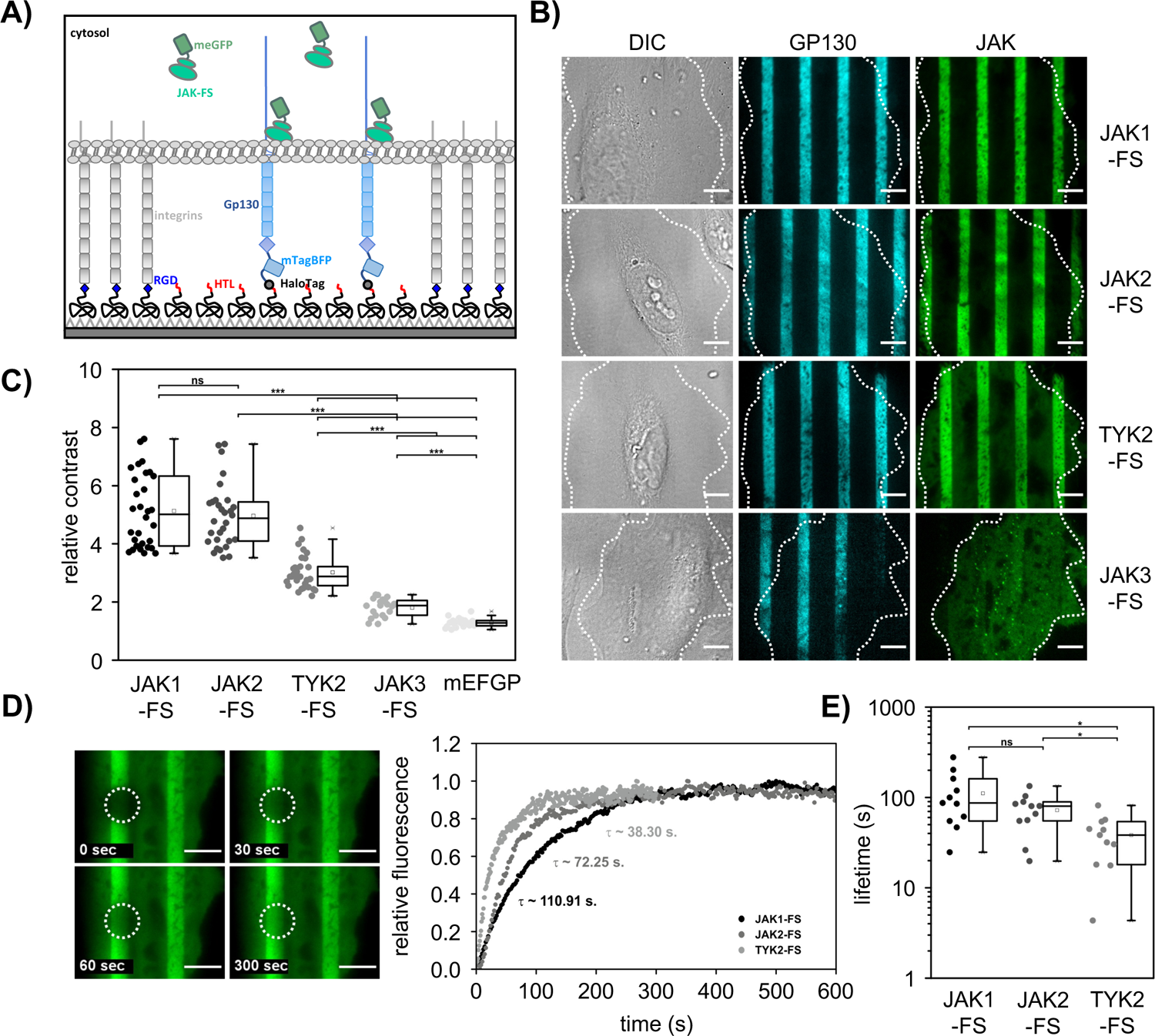
JAK1 and JAK2 bind Gp130 with similar affinities. **A)** Cartoon of live cell micropatterning to probe JAK binding to Gp130 in the plasma membrane. **B)** Representative experiments with micropatterned GP130 (cyan) interacting with the mEGFP-tagged FERM/SH2 domain of different JAKs (green). Scale bar: 10 µm. **C)** Comparison of the contrast in Gp130 micropatterns observed for different JAK FERM/SH2 domains. Each dot corresponds to an individual micropatterned cell. **D, F)** Gp130-JAK interaction stability probed by fluorescence recovery after photobleaching. **D**) Typical recovery experiment for the JAK1 FERM/SH2 domain (left) and representative recovery curves for JAK1-FS, JAK2-FS and TYK2-FS domains (right). Scale bar: 10 µm. **E**) comparison of the complex lifetimes obtained by fitting the recovery curve with a single exponential. Boxplots indicate the data distribution of second and third quartile (box), median (line), mean (square), 1.5× IQR (whiskers), and minimum/maximum (x). Each data point in C) and E) represents the analysis from one cell. Statistical analysis by kolmogorov smirnov test. Significances are indicated by asterisks (ns: p > 0.05, ∗: p ≤ 0.05, ∗∗: p ≤ 0.01, ∗∗∗p ≤ 0.001).

To obtain a more dynamic view of the Gp130-JAK interaction, we further analyzed the stability of JAK-Gp130 complexes using fluorescence recovery after photobleaching (FRAP). Since micropatterned Gp130 is immobilized in the micropatterns, the recovery curve reflects the exchange kinetics of photobleached JAK-FS with cytosolic, unbleached JAK-FS (Fig. 4D), which is typically dominated by the dissociation of the Gp130/JAK complex. By fitting the recovery curve with a monoexponential function, the lifetime of JAK-Gp130 complexes was determined (Fig. 4E). These analyses confirmed very similar complex stability for JAK1 and JAK2, and significantly reduced stability of TYK2 by a factor of 2-3 as compared to JAK1. For JAK3, this assay was not possible as the contrast was too low. These analyses confirmed that Gp130 can efficiently recruit JAK1 and JAK2, and to a somewhat lesser extent, also TYK2, but not JAK3. Since we observed prominent differences in Gp130 signaling even upon overexpression of JAK1, JAK2 and TYK, these cannot be explained by the relatively minor differences in receptor binding.

### The identity of the Gp130-JAK complex defines pSTAT activation by IL-6 in mammalian cells

Our studies in S2 cells suggest that the identity of the Gp130-JAK complex contribute to define pSTATs activation profiles by IL-6. However, whether these observations hold true in mammalian cells, where other cytokine regulatory mechanisms are in place remains unclear. To address this question, we explored how the identity of the Gp130/JAK complex impacts pSTAT activation in HeLa cells. To control specific JAK association to endogenous Gp130 (Fig. 5A), we used wt HeLa cells and a previously reported HeLa JAK1 KO clone (*36*). By combining these cells with siRNA targeting specific JAKs mRNAs, we generated JAK-specific HeLa cells (Fig. 5A). To generate JAK1-specific HeLa cells, wt HeLa cells were transfected with siRNAs targeting JAK2 and TYK2 for 48 hours. Knock down efficiencies of up to 90% for the two kinases were achieved (Fig. 5B). To generate JAK2- or TYK2-specific HeLa cells, JAK1 KO HeLa cells were transfected with siRNA targeting TYK2 or JAK2 respectively. Knock down efficiencies of up 90% were achieved in both cases (Fig. 5B). Leveraging this comprehensive control of JAK expression levels, we explored how the identity of the Gp130-JAK complex impacts pSTAT activation patterns in mammalian cells. JAK-specific HeLa cells were stimulated with HypIL-6 for 15 min and pSTAT1-Tyr701 and pSTAT3-Tyr705 levels were analyzed via phospho-flow cytometry (Fig. 5C). HeLa wt cells were used as a control of maximal HypIL-6 signaling. In response to HypIL-6 stimulation, JAK1-specific HeLa cells achieved pSTAT1 and pSTAT3 levels comparable to those induced in HeLa cells, suggesting that JAK1 is sufficient to induce maximal IL-6 signaling in these cells (Fig. 5C). However, JAK2- and TYK2-specific HeLa cells only triggered minimal amounts of STAT1 and STAT3 phosphorylation in response to HypIL-6 stimulation, with pSTAT1 being more affected that pSTAT3 (Fig. 5C). Interestingly, these observations are in stark contrast to our results in S2 cells, where Gp130-JAK2 and Gp130-TYK2 complexes triggered potent STAT3 phosphorylation in response to HypIL-6 stimulation. These results suggest that HeLa cells may have regulatory mechanisms in place to ensure signaling only by Gp130-JAK1 complexes.

**Figure 5.**
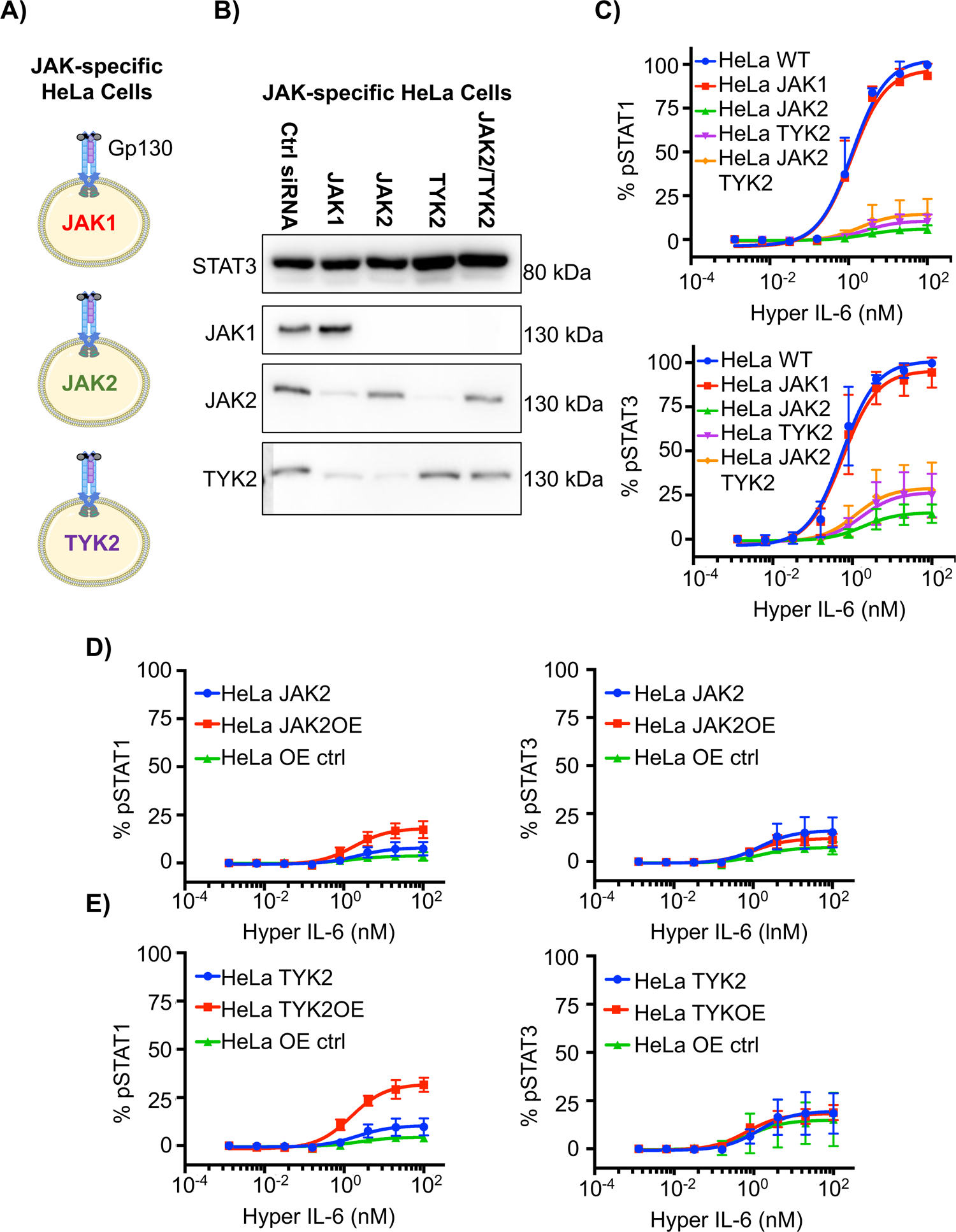
JAK1-, JAK2- and TYK2-Gp130 complexes trigger different profiles of STAT activation in HeLa cells. **A)** Cartoon highlighting the generation of JAK-specific HeLa cells. **B)** Western blot showing JAK1, JAK2 and TYK2 expression in HeLa cells transfected with the different siRNAs constructs. Total levels of STAT3 were used as a loading control. The siRNAs used specifically knock down their intended JAKs, with not off-target effect in the other JAK members. The western blot presented is a representative example of three independent experiments. **C)** Dose-response experiments of HeLa cells transfected with the indicated conditions. Transfected HeLa cells were stimulated with HypIL-6 for 15 min and levels of pSTAT1/pSTAT3 were measured via phosphoflow cytometry. Data are mean + /- SEM from three independent experiments. **D-E)** Dose-response experiments of HeLa cells transfected with the indicated conditions. HeLa JAK1 KO cells were treated with siRNAs targeting JAK2 (**D**) or TYK2 (**E**) and then transfected with TYK2 or JAK2 respectively. GFP-EPOR was used as an overexpression (OE) control. Transfected HeLa cells were stimulated with HypIL-6 for 15 min and levels of pSTAT1/pSTAT3 were measured via phosphoflow cytometry. Data are mean + /- SEM from three independent experiments.

A factor to consider when interpreting these results is that HeLa cells express very different levels of JAK1, JAK2 and TYK2. While HeLa cells express 209K copies of JAK1, they only express 6K copies of TYK2 and non-detectable levels of JAK2 (*37*). The functional consequence of this is that the number of Gp130-JAK complexes is significantly higher in a JAK1-specific HeLa, than in JAK2- or TYK2-specific HeLa cells. This in turn could result in the low pSTAT1/pSTAT3 levels induced by HypIL-6 in these cells. To account for this limitation, we increased the levels of JAK2 and TYK2 in HeLa cells to ensure that the number of Gp130/JAK complexes remain comparable in all instances. In a first step, JAK1 KO HeLa cells were transfected with JAK2 and TYK2 siRNAs to generate JAK-specific HeLa cells. Next, we overexpressed JAK2 or TYK2 in JAK2- or TYK2-specific cells respectively. JAK2- or TYK2-specific HeLa cells expressing high levels of JAK2 or

TYK2 were generated (Fig. S5A). Control experiments with truncated JAK2 variants established efficient competition with endogenous JAKs under these conditions (Fig. S5B). HeLa cells transfected with GFP-EPOR served as an overexpression control. Interestingly, overexpression of JAK2 only led to marginal increase in pSTAT1 levels, with no changes in pSTAT3 levels in response to HypIL-6 stimulation (Fig. 5D). By contrast, overexpression of TYK2 resulted in a 4-fold increase in pSTAT1 levels, but not pSTAT3 levels in response to HypIL-6 stimulation (Fig. 5E). In both instances, the overexpression of either JAK2 or TYK2 could not rescue IL-6 signaling to levels comparable to those induced by Gp130-JAK1 complexes. Collectively, our results showed that the low signaling activities induced by HypIL-6 in JAK2- and TYK2-specific HeLa cells is not a result of limited expression levels of these kinases, but to fundamental differences in the Gp130-JAK complexes engaged. These striking differences suggest that cells can rapidly adjust their responses to IL-6, simply by altering their JAK expression patterns.

### The overall number of Gp130 signaling complexes depend on JAK identity

We therefore hypothesized that JAKs may differentially affect receptor assembly. We have previously observed that JAKs can stabilize receptor dimerization at the plasma membrane, which can be ascribed to JAK-JAK interactions between the pseudokinase domains (*38–40*). Such interactions could depend on JAK identity and/or structural constraints imposed by binding to Gp130. To test whether JAKs affect the assembly of Gp130 signaling complexes in cells, we quantified receptor dimerization in the plasma membrane by single molecule co-tracking. To this end, Gp130 fused to an N-terminal ALFA-tag (*41*) was co-expressed with different JAKs fused to mEGFP in HeLa Gp130/JAK1 dKO cells. Cell surface-selective labeling of Gp130 was achieved with a mixture of anti-ALFA-tag nanobodies conjugated to two different, spectrally separable fluorophores (Fig. 6A). Formation of Gp130 dimers in the plasma membrane was detected and quantified by dual-color single molecule TIRF microscopy and co-tracking analysis (*42*). Strong dimerization of Gp130 induced by saturating concentrations of HypIL-6 was observed (Fig. 6B, C, S6A), with dimerization levels in line with previous studies using endogenous JAK1 (*29*). The dimerization levels observed in the presence of JAK2 and TYK2 were even somewhat higher as compared to JAK1. In the presence of JAK2 and TYK2, though, Gp130 already showed significantly increased dimerization levels in the absence of ligand (Fig. 6C). Likewise, diffusion properties of Gp130 were similar for different JAKs, with a minor decrease in the diffusion constant upon stimulation with HypIL-6 (Fig. S6C). In the presence of JAK-FS, similar HypIL-6-induced dimerization levels of Gp130 were observed, suggesting that the strong interaction of HypIL-6 dominates dimerization (Fig. S6E). By contrast, ligand-independent dimerization by JAK2 and TYK2 was largely abrogated, but recovered upon expression of JAKs only lacking the tyrosine kinase domain (JAK-ΔTK, Fig. S6D). These results confirmed weak interactions between the pseudokinase domains, which were surprisingly enhanced for JAK2 and TYK2 as compared to JAK1. We therefore concluded from these experiments that the reduced activity exerted by JAK2 and TYK2 is not related to compromised JAK-JAK interactions in the corresponding Gp130 signaling complexes.

**Figure 6.**
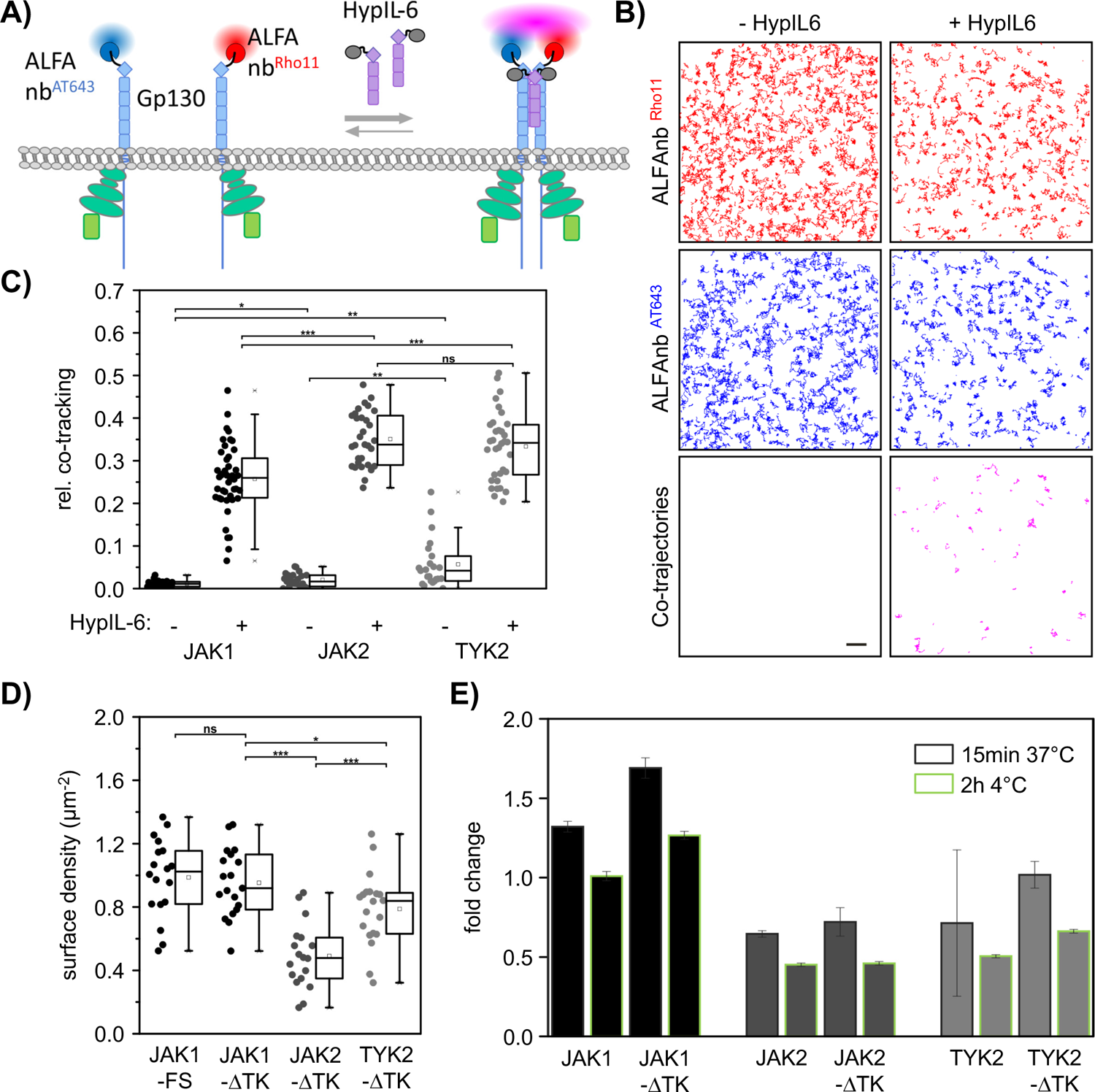
JAKs differentially affect formation of Gp130 signaling complexes. **A)** Cartoon depicting quantitative analysis of Gp130 dimerization by dual-color single molecule co-tracking. **B)** Representative single molecule trajectories and co-trajectories for Gp130-JAK1 before and after stimulation with HypIL-6. Scale bar: 2 µm. **C)** Comparison of HypIL-6 induced Gp130 dimerization upon co-expression of different JAKs performed in HeLa Gp130/JAK1 dKO cells. **D)** Single molecule localization density of fluorescent HypIL-6 bound to endogenous Gp130 at the surface of HeLa JAK1 KO cells expressing different JAKs. Boxplots indicate the data distribution of second and third quartile (box), median (line), mean (square), 1.5× IQR (whiskers), and minimum/maximum (x). Each data point in C) and D) represents the analysis from one cell. Statistical analysis by unpaired student’s t test. Significances are indicated by asterisks (ns: p > 0.05, ∗: p ≤ 0.05, ∗∗: p ≤ 0.01, ∗∗∗p ≤ 0.001). **E)** Overall ligand binding capacity of HeLa JAK1 KO cells expressing different JAKs quantified by flow cytometry.

We also explored dimerization of endogenous Gp130 in HeLa JAK1 KO cells by using a mixture of site-specifically fluorescent-labeled labeled HypIL-6 (Fig. S6B). To exclude bias by activation of downstream signaling, JAK-ΔTK variants were used for complementing with different JAKs. Surprisingly, we observed strongly reduced cell surface levels of HypIL-6 for cells expressing JAK2 as compared to JAK1 and TYK2 (Fig. 6D). With a total concentration of 10 nM HypIL-6 applied in these experiments, we expect saturation of all receptor binding sites, and therefore interpret these differences being caused by changes in Gp130 cell surface expression. We therefore explored this effect more systematically by quantifying total HypIL-6 binding by flow cytometry. Strikingly, differential binding capacities of endogenous Gp130 depending on different JAKs was confirmed, with JAK2 and TYK2 yielding ∼50% of the binding levels obtained for JAK1 (Fig. 6E). While JAK1 expression enhanced the number of HypIL-6 binding sites as compared to the basal level, these where strongly reduced upon expression of JAK2 and TYK2. This could be related to the stronger basal dimerization of Gp130 by JAK2 and TYK2 as compared to JAK1. Interestingly though, significant differences in cell surface expression could already be observed upon co-expression of JAK-FS fragments, yet less pronounced as in the presence of the full-length JAK. Strongest differences were found in experiments carried out at 4°C, which should minimize uptake by endocytosis. These results suggest that JAK identity strongly contributes to Gp130 cell surface expression, which is partly encoded by the JAK-FS domain, but possibly also involves interactions between JAKs in the Gp130 dimer and activation of downstream signaling.

To pinpoint whether differential modulation of Gp130 expression by JAK1, JAK2 and TYK2 is responsible for the strongly altered pSTAT patterns observed in mammalian cells, we performed activity with ectopically expressed Gp130 in combination with different JAKs. Upon gating for similar Gp130 expression levels (Fig. S7), very similar levels of pSTAT1 activation were observed for all three JAKs (Fig. S7). These results corroborate that the substantial differential activity observed in HeLa cells can be ascribed to Gp130 trafficking depending on JAK identity.

## DISCUSSION

In this study we have used *Drosophila* S2 cells and engineered HeLa cells to investigate how Gp130 translates cytokine binding events into activation of specific intracellular signaling networks and responses. Three main finding arise from our study: (1) The identity of the Gp130-JAK complex defines the profile of STAT activation induced by IL-6. The JAK member associated with Gp130 defines the pool of phosphotyrosines available in Gp130 for STAT recruitment and thus the identity and potency of the STAT molecules engaged. (2) The different enzymatic activities elicited by the Gp130-JAK complexes cannot be assigned to differences in binding affinity between Gp130 and JAK1, JAK2 or TYK2 or defects in IL-6-Gp130 hexameric complex formation. This suggest that Gp130 may implement topological constrains in JAK binding to ensure correct signaling only in the presence of the right JAK protein. (3) Gp130 cell surface expression strongly depends on JAK identity, despite similar binding affinities. In summary, our study describes a new layer of cytokine signaling regulation, whereby Gp130-JAKs complexes exhibit differential functional coupling that contribute to expand Gp130 signaling plasticity.

One question that arises from our study is why Gp130 needs to bind diverse JAKs? Although this is not a unique feature of Gp130, other receptors such as TpoR, OSMR and IL12Rβ2 have also been described to bind multiple JAKs (*18, 43, 44*), the vast majority of cytokine receptors show a strong specificity for a given JAK protein. Interestingly, all cytokine receptors binding multiple JAKs are ancestral receptors which origin in some instances, e.g. Gp130, can be traced back to Drosophila (*45*). Thus, it is possible that JAK binding specificity only evolved at later stages, while JAK binding promiscuity further enhanced signalling plasticity. Our study suggests that in mammals, the ability of Gp130 to bind different JAKs allows this receptor to fine-tune its responses by switching the JAK used, ultimately impacting the pool of phosphotyrosines available in its intracellular domain for STAT binding. In agreement with this model, a recent study reports the ability to decouple inflammation from tissue regeneration activities downstream of Gp130 by altering the number of Tyrosines in Gp130 available for phosphorylation (*46*).

Our data clearly demonstrate that JAK1, JAK2 and to a lower extent TYK2 bind with comparable affinities to Gp130. Thus, the number of Gp130-JAK1, Gp130-JAK2 and Gp130-TYK2 complexes in each cell subset will be determined by the different absolute and relative concentrations of those JAKs in each cell. Recent studies have shown that JAK concentrations dramatically change not only between cell subsets, but also depending on differentiation state of any cells (*47*). Thus, it is tempting to speculate that cells can change their JAK concentrations in response to environmental cues to fine-tune their responses to cytokines. Since different Gp130-JAK complexes elicit very different signaling outputs, deregulation of this equilibrium could lead to disease. In line with this model, several reports have described that tumour cells can use the IL-6/Gp130/JAK2/STAT3 pathway to drive cancer progression and resistance. Breast cancer cells preferentially use the IL-6/JAK2/STAT3 pathway to promote growth and survival by maintaining a stem cell-like cancer cell population (*48*). IL-6 expression and activation of JAK2/STAT3 signaling pathway contribute to poor prognosis and reduced survival in nasopharyngeal carcinoma patients (*49*). Increased TrkC expression promotes tumorigenicity, metastasis, and self-renewal of metastatic breast cancer by regulating the IL-6/Gp130/JAK2/STAT3 axis (*50*). Although the contribution of JAK1 to the observed phenotype was not explored in these studies, their main conclusions agreed with our observations in S2 cells where Gp130-JAK2 complexes exhibited a strong bias towards STAT3 activation. Under normal conditions IL-6 induces activation of both STAT1 and STAT3 proteins (*15*). However, in certain cancers, where JAK2 expression may be favoured (*50*), IL-6 could preferentially engage Gp130-JAK2 complexes that would exhibit a bias towards STAT3 activation, leading to strong tumour proliferation and survival.

Although JAK1 and JAK2 bind with very similar affinities to Gp130, their activities profiles in HeLa cells are strikingly different. This was not the result of lower JAK2 expression in HeLa cells since over-expression of this kinase failed to rescue signaling by Gp130-JAK2 complexes. How do Gp130-JAK1 and Gp130-JAK2 elicit such different signaling outputs? JAKs bind two membrane proximal regions, known as Box1 and Box2, in the cytokine receptor intracellular domains via their FERM-SH2 domain (*51*). Recent structural studies have provided insights into how JAKs bind this Box1/Box2 region, revealing key binding determinants that contribute to receptor-JAK specificity (*39, 51, 52*). One example is found in the positioning of the F2-a3 helix in different JAKs, which we have recently proposed interact with the membrane and thus stabilize JAK orientation in the signalling complex (*38*). This helix is rotated in JAK2, as compared to JAK1 and TYK2, which may have implication for the orientation at the membrane, but may also explain altered trafficking (s. below). However, we still lack structural information regarding how a cytokine receptor can bind different JAKs, limiting our ability to provide molecular insights into how these interactions regulate cytokine responses. Interestingly, cytokine receptors that bind multiple JAKs appear to have a conserved Box2 PxP motif, which critically contributes to JAK binding (*51, 52*), and relatively highly conserved recognition by the JAK’s FERM/SH2 domains has been observed. However, differential activities of the thrombopoietin receptor bound to different JAKs has been related to differences in orientation (*53, 54*). These observations suggest that JAKs members may differ in the structural organization of pseudokinase/kinase module relative to the FERM domain, in line with relatively high diversity on the connecting linker. This model agrees with a recent study that suggests that the topology of the JAK-receptor complex critically contributes to weak signal activation by type III interferons (*55*). Moreover, receptor associated JAKs require a complex inter-relationship for efficient signal activation (*35, 38*). Small topological alteration in this JAK-receptor interaction could have dramatic consequences for signaling, which may explain the differences in pSTAT signatures observed in S2 cells.

However, our data show that, despite similar binding affinity and stability, Gp130 surface levels are strongly affected by the identity of the JAK molecule bound to this receptor, with JAK2 inducing a strong decrease in Gp130 surface expression in mammalian cells. Beyond their Tyr kinase activity, JAKs also play a chaperone role regulating the surface expression of some cytokine receptors (*18, 31, 56-58*). JAKs mask a three dileucine-like motif within the inter-Box-1/2 region, which contributes to destabilizing the cytokine receptor in the absence of JAKs (*18, 31, 56-58*). This motif can be found in Gp130, but regulates ligand-induced Gp130 internalization rather than Gp130 levels at homeostasis (*59*). Indeed, JAK1 KO HeLa cells express comparable levels of Gp130 as compared to WT HeLa cells (Fig. 6E), suggesting that the lower levels of Gp130 found in the context of JAK2 and TYK2 expression do not result from destabilization of the Gp130 dileucine-like motif. JAKs also play an active role in defining intracellular trafficking of cytokine-receptor complexes. JAKs can interact with the endosomal machinery and contribute to sort the cytokine-receptor complex to define compartments where they initiate defined signaling programs (*60*). Thus, Gp130-JAK1 complexes could follow a different endosomal path than Gp130-JAK2 or Gp130-TYK2 complexes, which ultimately could impact their signal activation potency. Addressing the endosomal routes followed by different GP130-JAK complex would thus be crucial to better understand Gp130 biology and inform future engineering approaches to fine-tune Gp130 signaling. Overall, our work defines an intricate, multi-tiered functional coupling between Gp130 and JAKs that contribute to expand the bioactivity range of this crucial shared receptor. Even more complexity can be expected by formation of GP130 dimers with heteromeric JAK combinations. With Gp130 being involved in diverse cytokine-receptor complexes, broad and complex implications of our findings can be expected. Understanding how the different Gp130-JAK complexes regulate Gp130 biology in different physiological conditions and disease will prove crucial to untangle Gp130 complex biology and design more efficient Gp130-based therapies.

---

**Figure S1. Setting up the transfection of the IL-6-Gp130 signaling network in S2 cells. A)** S2 cells were nucleofected with a fix concentration GFP-Gp130, STAT3 cDNA and different concentrations of JAK1 cDNA and then stimulated with HypIL-6 (20 nM) for 15 minutes or left untreated. Amounts of JAK1 DNA greater than 3 µg resulted in a significantly higher pSTAT3-Tyr705 signal in the absence of stimulation. Data are mean + /- SEM from three independent experiments. **B)** S2 cells were nucleofected with GFP-Gp130/JAK1/STAT3 and stimulated with HypIL-6 for up to 180 minutes. pJAK1 and pSTAT3 levels were assessed by western blotting. Total STAT3 and tubulin were used as a loading control. Non-transfected S2 cells were included as comparison. The western blot presented is a representative example of three independent experiments. **C)** S2 cells were nucleofected with GFP-Gp130, JAK1 and STAT3 and then stimulated with 20 nM of HypIL-6 for 0-210 minutes. After 15 minutes of stimulation, 2 μM Tofacitinib was added which resulted in a rapid decrease of pSTAT3-Tyr705 signal. Data are mean + /- SEM from three independent experiments.

**Figure S2. JAK/STAT activation profiles induced by IL-6 in S2 cells.** Dose-response profiles induced by HypIL-6 stimulation in S2 cells transfected with each JAK (JAK1, JAK2, JAK3 and TYK2) and STAT (STAT1-6). Data was normalized to the JAK1 condition. Data are mean + /- SEM from three independent experiments.

**Figure S3. S2 cells can express all components of the JAK/STAT pathway.** Western blots showing the expression of all components of the JAK/STAT pathway in transfected S2 cells used during the dose-response studies illustrated in Fig. S2. The western blot presented is a representative example of three independent experiments.

**Figure S4. STAT activation profiles induced by Gp130 mutants in S2 cells. A)** Dose-response profiles induced by HypIL-6 stimulation in S2 cells transfected with the indicated Gp130 mutants. Data was normalized to the JAK1 condition. Data are mean + /- SEM from three independent experiments. **B)** Dot plot showing pSTAT1/pSTAT3 ratio induced by HypIL-6 in S2 cells transfected with wt Gp130 or Gp130 3F mutant. Data are mean + /- SEM from three independent experiments. **C)** Representative FACS plot showing Gp130 transfection efficiencies in S2 cells. The truncated Gp130 mutant exhibit higher levels of expression than Gp130 wt. **D)** Dose-response profiles induced by HypIL-6 stimulation in S2 cells transfected with the truncated Gp130 mutant. Data was normalized to the JAK1 condition. Data are mean + /- SEM from three independent experiments.

**Figure S5. JAK/STAT activation profiles induced by IL-6 in HeLa cells. A)** Western blots showing the expression of JAK2 and TYK2 in transfected HeLa cells. The western blot presented is a representative example of three independent experiments. **B**) STAT3 phosphorylation in HypIL-6-stimulated wt HeLa cells expressing inactive JAK-ΔTK fused to mEGFP as quantified by phospho-flow cytometry. Cells were gated for different expression levels of JAK-ΔTK as indicated on the top of the graphs. Data are mean + /- SEM from one experiment. **C**) Relative pSTAT1 and pSTAT3 levels induced by the kinase domains of JAK1, JAK2 and TYK2 expressed HeLa cells.

**Figure S6. Gp130 dimerization in presence of JAK overexpression probed by single molecule co-localization and co-tracking TIRF microscopy. A)** Surface density of Gp130 ectopically expressed in HeLa Gp130/JAK1 dKO cells. **B)** Cartoon depicting labeling and quantification of endogenous Gp130 in HeLa JAK1 KO cells by DY547P1 and DY647P1 conjugated HypIL-6. Ectopic expression of GFP-tagged JAKs was checked for each recorded cell. **C)** Average diffusion constants of cell surface Gp130 in HeLa Gp130/JAK1 dKO cells overexpressing one of the JAKs. **D)** Gp130 homo-dimerization in HeLa Gp130/JAK1 dKO expressing one of the JAK-ΔTK with and without HypIL-6 incubation. **E)** Gp130 homo-dimerization in HeLa Gp130/JAK1 dKO expressing one of the JAK-FS with and without HypIL-6 incubation. Boxplots indicate the data distribution of second and third quartile (box), median (line), mean (square), 1.5× IQR (whiskers), and minimum/maximum (x). Each data point in A), C), D), and E) represents the analysis from one cell. Statistical analysis by unpaired student’s t test. Significances are indicated by asterisks (ns: p > 0.05, ∗: p ≤ 0.05, ∗∗: p ≤ 0.01, ∗∗∗p ≤ 0.001).

**Figure S7. Robust signal activation by all JAKs upon ectopic expression of Gp130. A)** STAT1 phosphorylation in HeLa JAK1 KO cells transfected with different JAKs. pSTAT1 was quantified by phospho-flow cytometry without agonist and after stimulation with 10 nM HypIL-6 for 15 min. Non-transfected cells were used as control (empty boxes). **B)** Same experiment carried out with additional transfection of Gp130 and gating for similar receptor expression levels.

## MATHERIAL AND METHODS

### Cells and Media

All cells were handled in sterile laminar flow culture hoods. Mammalian cells were cultured in incubators which were maintained at 37DC, 5% C02. The insect cells Trichoplusia ni (Hi5) (Thermo Fisher Scientific) and Spodoptera frugiperda (SF9) (Thermo Fisher Scientific) were cultured with gentle agitation (120 rpm) at 27DC. Drosophila melanogaster cells (S2) were cultured without agitation at 27DC.

All cells were cultured in the following media and buffers:

| Solution | Composition |
| --- | --- |
| ***Drosophila* Schneiders Media** | Schneider's *Drosophila* Medium (Biosera)<br>10% heat-inactivated fetal bovine serum (HI-FBS) (Gibco)<br>100 IU/mL Penicillin and 100 µg/mL Streptomycin (Gibco).<br>Filter Sterilized |
| **Complete RPMI (cRPMI)** | RPMI-1640 (Gibco)<br>2 mM L-glutamine (Gibco)<br>10% foetal bovine serum (FBS) (Gibco),<br>100 IU/mL Penicillin and 100 µg/mL Streptomycin (Gibco). |
| **Hi5 media** | Lonza Insect-Xpress with glutamine (SLS)<br>Gentamicin (10 µg/mL) (Gibco). |
| **SF9 media** | SF-900™ III serum free medium (SFM) (Gibco)<br>10 µg/mL Gentamicin (Gibco). |
| **PBSA** | Phosphate buffered saline, pH 7.2 (made in-house by SLS Central Technical Services, University of Dundee)<br>0.5% Bovine Serum Albumin (BSA) (Sigma) |
| **PBE** | Phosphate buffered saline (PBS), pH 7.2 (made in-house by SLS Central Technical Services, University of Dundee)<br>0.5% BSA (Sigma)<br>2 mM EDTA (SLS Central Technical Services) |
| **Complete DMEM** | DMEM (Gibco)<br>2 mM L-glutamine (Gibco)<br>10% foetal bovine serum (FBS) (Gibco),<br>100 IU/mL Penicillin and 100 µg/mL Streptomycin (Gibco). |

### Nucleofection of S2 cells

For each nucleofection, ∼3.5×10^6^ S2 cells were harvested and pelleted by centrifugation at 100 g for 5 minutes. The supernatant was thoroughly removed, and the cell pellet was resuspended in 100 µl of the in-house nucleofection buffer. The following amounts of DNA were added after optimization levels were obtained: GFP-Gp130 pAc5.1 (5 µg), JAK1/JAK2/JAK3/TYK2 (3 µg) and STAT1-6 (5 µg). Each DNA construct was expressed within the pAc5.1 vector for expression within S2 cells. The nucleofection solution was thoroughly mixed and added to a generic electroporation cuvette (VWR, 2-mm gap). All S2 cells were nucleofected using the G-030 programme on the Amaxa Nucleofector^®^ II (Lonza). Immediately after nucleofection, 500 µl of Schneider’s *Drosophila* media (S2 media) was added to the cuvette and the cells were seeded into 6-well dishes containing 1.5 mL of complete S2 cell media, taking care to avoid bubbles. The nucleofected samples were then incubated at 27°C in the dark for 48 hours before being used in subsequent experiments.

### siRNA silencing

HeLa WT and HeLa JAK1 knock out (KO) cells were seeded at 2×10^5^ cells per well in a six well plate 12 hours before further treatment. After incubating, the cells were transfected with various siRNAs (Dharmacon) using DharmaFECT 1 transfection reagent (Dharmacon, Cat #T2001-02) following the manufacturer’s instructions. Cells were either transfected with JAK2 siRNA (all 4 pools pooled), TYK2 siRNA (all 4 pools pooled) or lastly both JAK2 and TYK2 siRNA in combination. After siRNA transfections, cells were incubated for 48 hours at 37LJC and subsequently prepared for immunoblotting analysis to check the level of gene silencing and phosphoflow cytometry to check signalling dynamics (pSTAT1-Tyr701/pSTAT3-Tyr705).

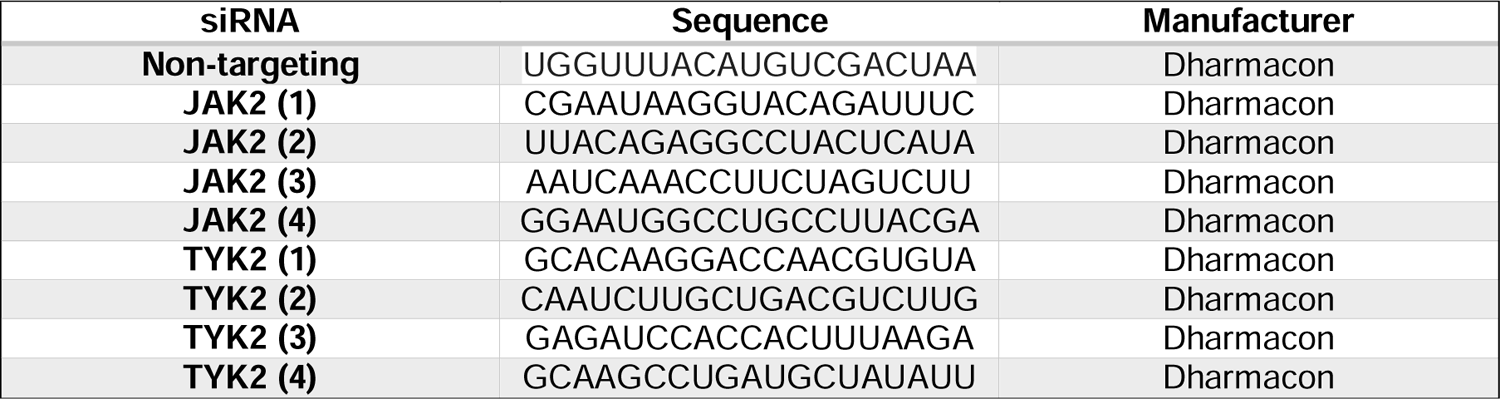

### HeLa cells transfections

HeLa WT and HeLa JAK1 KO cells were seeded at 2×10^5^ cells per well in a six well plate and transfected with siRNA 24 hours before PEI transfections. To transfect cells with PEI, DNA was diluted in incomplete DMEM. The optimized amount for these studies was 3 ug DNA in 200 µl of incomplete media. For HeLa cells, pSems constructs were used. 15 µl of PEI was added to the diluted DNA and the sample vortexed for 15 seconds. The transfection mix was incubated at room temperature for 20-30 minutes. Media was removed from cells in 6-well dishes and these washed with DPBS. 1 mL of incomplete DMEM was added to the transfection mix and the solution added dropwise to the appropriate pre-washed well. Cells were incubated for 3 hours at 37LJC. After incubation, a further 2 mL of complete DMEM was added to the wells and the cells incubated for a further 24 hours at 37LJC. Subsequently, cells were prepared for immunoblotting analysis to check the level of gene expression and phosphoflow cytometry to check signalling dynamics (pSTAT1-Tyr701/pSTAT3-Tyr705).

### Flow cytometry and phospho-flow studies

All antibodies, clones and suppliers are listed below:

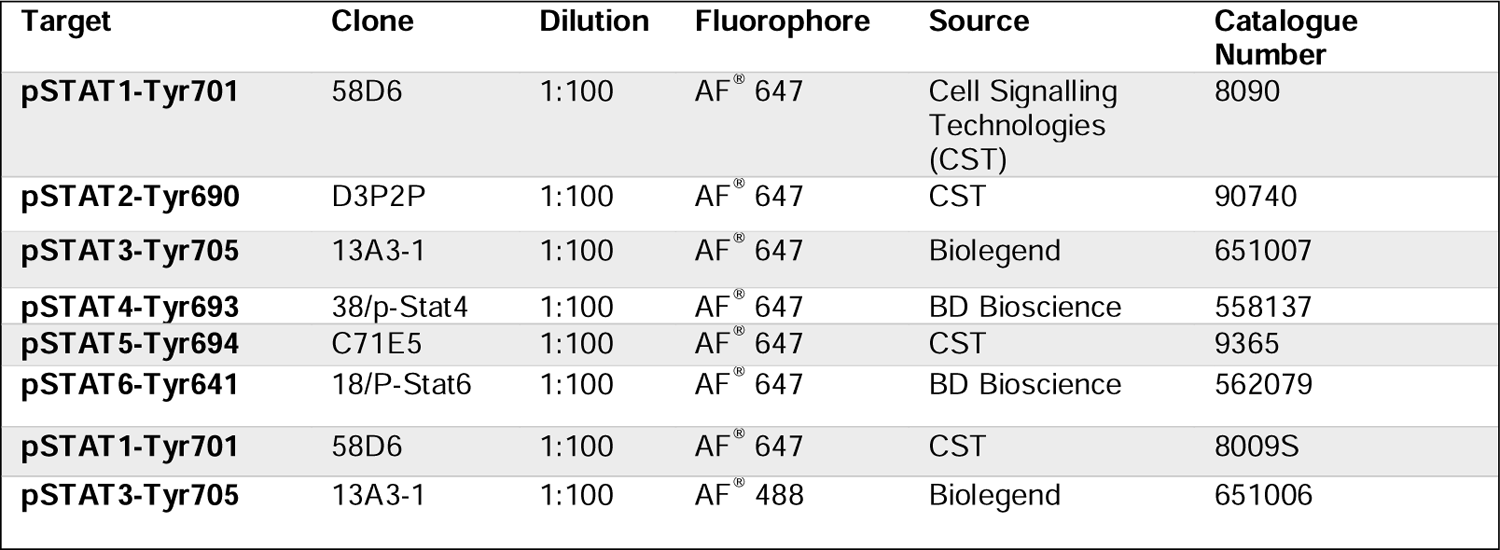

| Target | Clone | Dilution | Fluorophore | Source | Catalogue Number |
| --- | --- | --- | --- | --- | --- |
| pSTAT1-Tyr701 | 58D6 | 1:100 | AF<sup>®</sup> 647 | Cell Signalling Technologies (CST) | 8090 |
| pSTAT2-Tyr690 | D3P2P | 1:100 | AF<sup>®</sup> 647 | CST | 90740 |
| pSTAT3-Tyr705 | 13A3-1 | 1:100 | AF<sup>®</sup> 647 | Biolegend | 651007 |
| pSTAT4-Tyr693 | 38/p-Stat4 | 1:100 | AF<sup>®</sup> 647 | BD Bioscience | 558137 |
| pSTAT5-Tyr694 | C71E5 | 1:100 | AF<sup>®</sup> 647 | CST | 9365 |
| pSTAT6-Tyr641 | 18/P-Stat6 | 1:100 | AF<sup>®</sup> 647 | BD Bioscience | 562079 |
| pSTAT1-Tyr701 | 58D6 | 1:100 | AF<sup>®</sup> 647 | CST | 8009S |
| pSTAT3-Tyr705 | 13A3-1 | 1:100 | AF<sup>®</sup> 488 | Biolegend | 651006 |

96-well plates were prepared with 50µl of cell suspensions at 2×10^5^ cells/ml/well. Cells were stimulated with a set of different concentrations to obtain dose-response curves. To this end cells were stimulated for 15 min at 37°C with HypIL-6 followed by PFA fixation (2%) for 15 min at RT. After fixation, cells were spun down at 300g for 6 min at 4°C. Cell pellets were resuspended and permeabilized in ice-cold methanol and kept for 30 min on ice. After permeabilization cells were fluorescently barcoded according to Krutzil et al., 2001 (*61*). In brief: using two NHS-dyes (PacificBlue, #10163, DyLight800, #46421, Thermo Scientific), individual wells were stained with a combination of different concentrations of these dyes. After barcoding, cells are pooled and stained with the indicated pSTATs antibodies at a 1:100 dilution in PBS+0.5%BSA for 1h at RT. Cells were analzyed on a flow cytometer (Beckman Coulter, Cytoflex S) and individual cell populations were identified by their barcoding pattern. Mean fluorescence intensity (MFI) of pSTAT was measured for all individual cell populations.

### Western blotting

2×10^6^ transfected S2 cells were lysed in 50 μl of RIPA buffer (#FNN0021, Thermo Fisher Scientific) and incubated on ice for 10 minutes. Samples were spun at 15000 rpm for 10 minutes at 4°C and the supernatants retained. As a next step, 6X reducing buffer was added to the lysate supernatants and samples heated at 80°C for 10 minutes. Denatured samples were then run out on 7% polyacrylamide gels then transferred to nitrocellulose membranes using a Western blotting transfer apparatus, used in accordance with the manufacturer’s instructions (BioRad). Membranes were incubated for 1 hour in 5% w/v BSA in PBST (1xPBS, 0.1% Tween 20). The membranes were washed three times in PBST for 5 minutes and incubated in primary antibodies for 3 hours at room temperature or 12 hours at 4°C. Membranes were washed three times for 5 minutes each in PBST and incubated with a 1:5000 dilution of horseradish peroxidase-conjugated anti-mouse or anti-rabbit antibodies (Peroxidase-AffiniPure Donkey anti-Rabbit IgG-711-035-152, Peroxidase-AffiniPure Donkey anti-Mouse IgG-715-035-150, Stratech) for 1 hour at room temperature. Blots were then washed in PBST four times for 15 minutes each before being developed using Immobilon^TM^ Western Chemiluminescent HRP Substrate in accordance with manufacturer’s instructions (Millipore).

Antibody list for Western Blotting:

| Protein | Epitope | Concentration | Source | Catalogue No. |
| --- | --- | --- | --- | --- |
| **Gp130** | Total | 1:1000 | Santa Cruz Biotechnology | Sc-376280 |
| **STAT1** | Total | 1:1000 | Cell Signalling Technology (CST) | 9172 |
| **STAT2** | Total | 1:1000 | CST | 72604 |
| **STAT3** | Total | 1:1000 | CST | 9132 |
| **STAT4** | Total | 1:1000 | CST | 2653 |
| **STAT5** | Total | 1:1000 | CST | 9363 |
| **STAT6** | Total | 1:1000 | CST | 9362 |
| **JAK1** | Total | 1:1000 | CST | 3344 |
| **JAK2** | Total | 1:1000 | CST | 3230 |
| **JAK3** | Total | 1:1000 | CST | 3775 |
| **TYK2** | Total | 1:1000 | CST | 9312 |
| **pJAK1** | Phospho-Tyr1034/35 | 1:1000 | CST | 74129 |
| **pSTAT3** | Phospho-Tyr705 | 1:1000 | CST | 9145 |
| **Tubulin** | Total | 1:50 | Developmental Studies Hybridoma Bank (DSHB) | AB_1157911 |
| **LaminC** | Total | 1:50 | DSHB | AB_528332 |
| **GAPDH** | Total | 1:1000 | CST | 2118 |

### Cell micropatterning and image analysis

Binding of JAKs to Gp130 in the plasma membrane was quantified by live cell micropatterning using the HaloTag (*62*) for capturing cell surface receptors (*34*). Surfaces functionalized with the micropatterned HaloTag ligand (HTL) were fabricated by microcontact printing as described in detail previously (*33*). Standard glass coverslips for fluorescence microscopy were cleaned in a plasma cleaner for 10 min. PDMS stamps were inked with 0.5 mg/mL poly-L-lysine-graft-poly (ethylene glycol) (PLL-PEG) conjugated with the HTL (PLL-PEG-HTL) in PBS buffer for 10 min and then placed with additional weight onto the glass coverslips for 10 min in order to generate HTL micropatterns. After removing the stamps, the coverslips were incubated with a 4:1 mixture of 0.5 mg/mL of methoxy-terminated PLL-PEG (PLL-PEG-MeO) and 0.5 mg/mL PLL-PEG conjugated with the peptide RGD (PLL-PEG-RGD) (*63*) in PBS buffer for 2 min to backfill the uncoated area to allow cell adhesion. The surface was then rinsed in MilliQ water and dried with nitrogen.

Cells were transiently transfected with gp130 N-terminally fused to the HaloTag and mTagBFP (HaloTag-mtagBFP-gp130), truncated JAK1/JAK2/TYK2 and JAK3 comprising only the FERM and SH2-like domains C-terminally fused to mEGFP (JAK-FS). Transfected cells were transferred onto micropatterned surfaces 24-36 h post transfection and cultured for 15-20 h with medium containing penicillin and streptomycin (PAA). Micropatterned cells were then imaged by TIRF microscopy using an inverted microscope (Olympus IX81) equipped with a 4-line TIRF condenser (Olympus TIRF 4-Line LCl), a CMOS camera (ORCA-Flash 4.0, Hamamatsu) and lasers at 405 nm (100 mW), 488 nm (150 mW). A 60X and a 100X objective with a numerical aperture of 1.49 (UAPON 60×/1.49, Olympus, UAPON 100x/1.49) was used for TIRF excitation. The excitation beam was reflected into the objective by a quad-band dichroic mirror (zt405/488/561/640rpc) and the fluorescence was detected through a quadbandpass filter (BrightLine HC 446/523/500/677). For multicolor experiments, a fast emission filter wheel equipped with suitable emission filters (BrightLine HC 445/45, BrightLine HC 525/50, BrightLine HC 600/37 and BrightLine HC 697/58) was utilized to avoid spectral cross-talk. Data acquisition was performed with the acquisition software Olympus CellSens 2.2.

Image analysis and image processing were performed using ImageJ/ Fiji (NIH, Bethesda, MD). Image processing comprised cropping, scaling, rotation as well as adjustment of brightness and contrast levels. The fluorescence contrast of the bait proteins (*C_bait_*) was calculated as:

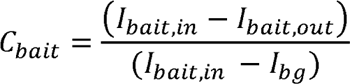

where *I_bait,in_* and *I_bait,out_* denote the mean pixel intensities from selected ROIs inside and outside, respectively, of the HTL-functionalized areas and *I_bg_* the background intensity from the glass surface obtained from a ROI outside the region covered by the cells. The contrast of the prey proteins *C_prey_* was accordingly obtained from the background corrected mean pixel intensities from selected ROIs inside and outside HTL-functionalized areas:

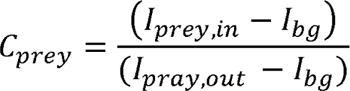

Since *C_prey_*is dependent on *C_bait_*, *C_prey_* was corrected for variances in *C_bait_* by dividing *C_prey_* by *C_bait_*:

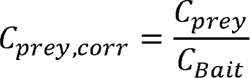

For statistical analysis, 30 cells were analyzed and the calculated contrast values were visualized in a box-plot. Two-sample Kolmogorov-Smirnov-Tests were performed to calculate statistical significances.

The interactions stability was determined by fluorescence recovery after photobleaching (FRAP). A rectangular region of interest (ROI) within the bleached area of the pattern and a rectangular ROI within the bleached area but outside the patterned area were chosen for obtaining intensity values per pixel over time, respectively. Corrected FRAP curves were determined using the following equation:

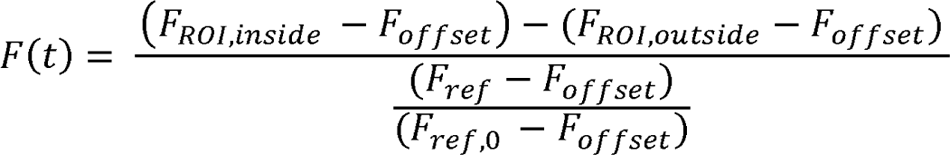

With *F_ROI,inside_* and *F_ROI,outside_*being the fluorescence intensities inside and outside the pattern, respectively, within the bleached spot. *F_ref_* is the fluorescence intensity of an unbleached ROI inside the micropattern, and *F_ref,0_*is the fluorescence intensity of this ROI before the bleaching experiment. *F_ref_*was implemented as a normalization factor to correct for photobleaching during FRAP experiments. The offset intensity (*F_offset_*) was determined from an ROI outside of the cell and was subtracted from all intensity values. Image analysis to obtain corrected FRAP curves was performed using a MATLAB script. In order to obtain lifetime values, the corrected FRAP curves *F(t)* were fitted by a monoexponential function.

### Surface expression by Flow Cytometry

HeLa cells were transiently transfected with either full-length JAK1, JAK2 or TYK2 C-terminally fused to meGFP (JAK1-meGFP, JAK2-meGFP, TYK2-meGFP) or truncated JAK comprising only the FERM and SH2-like domains (JAK-FS) or additionally the pseudokinase domain (JAK-ΔTK). To monitor endogenous gp130 surface expression, cells were incubated with 10 nM HypIL-6 labelled with Dy647 (^Dy647^HypIL-6) and incubated either at 37°C for 15 min or at 4°C for 2h. The cells were washed with PBS and afterwards detached by incubating with Accutase for 2 min at 37°C. Cells were centrifuged at 300 g for 5 min and the supernatant was discarded. After fixation with 4% PFA at RT for 15 min, cells were washed in PBS and centrifuged at 300 g for 5 min. The mean fluorescence intensity (MFI) of ^Dy647^HypIL-6 was measured by the BC CytoFLEX S flow cytometer and data analyzed with BC CytExpert software.

### Live-cell dual-color single-molecule imaging studies

Dual-color single-molecule imaging was carried out by total internal reflection fluorescence microscopy (TIRFM) using an inverted microscope (IX83-P2ZF, Olympus) equipped with a motorized quad-line TIR illumination condenser (cellTIRF-4-Line, Olympus) as recently described in detail (*42*). The fluorophores mEGFP, ATTO Rho11 and ATTO 643 were excited using a 100× oil immersion objective (UPLAPO100XOHR, NA 1.5, Olympus) at 488 nm (LuxX 488-200, max. 200 mW, Omicron), 561 nm (2RU-VFL-P-500-560-B1R, MPB Communications) and 642 nm (2RU-VFL-P-500-642-B1R, MPB Communications), respectively. Fluorescence was filtered by a penta-band polychroic mirror (zt405/488/561/640/730rpc, Semrock) and excitation light was blocked by a penta-band bandpass emission filter (BrightLine HC 440/521/607/694/809, Semrock). Up to four channels could be simultaneously acquired by using the four quadrants of a single back-illuminated sCMOs camera (Hamamatsu ORCA Fusion-BT, Hamamatsu Photonics) and a four-color image splitter (MultiSplit V2, CAIRN). The latter is equipped with three dichroic beamsplitters at 560 nm, 640 nm and 730 nm (H 560 LPXR-UF2, H 643 LPXR-UF2, AHF; T 725 LPXR-UF2, Chroma) and four single-band bandpass emission filters (BrightLine HC 520/35, BrightLine HC 600/37, BrightLine HC 809/81, Semrock; ET 685/50, Chroma). For dual channel imaging, only the orange (ATTO Rho11) and red (ATTO 643) channel were acquired. The green (mEGFP) channel was used to check for ectopic JAK-mEGFP expression. A 2×2 pixel binning was applied to obtain a pixel size of 130 nm. The focus was continuously stabilized during the experiment by a hardware autofocus-system (IX3-ZDC2, Olympus) using an internal laser diode at 830 nm.

For single molecule tracking and co-tracking, GP130 WT fused to an N-terminal ALFA-tag were employed for selective cell surface labelling via anti-ALFA nanobodies (*41*). For this purpose, the ORFs of GP130 WT lacking the signal peptide was cloned into the vector pSems comprising the signal peptide of Igκ chain followed by ALFA peptide and a short peptide linker (pSems-leader-ALFA-linker-GP130). GP130 KO HeLa cells (*29*) were transfected with pSems-leader-ALFA-linker-GP130 in 6 cm dishes at 70% confluency using a polyethylenimine (PEI) transfection protocol (*64*) on the day before imaging. For imaging transiently transfected cells were seeded on microscopy cover slides coated with a 50/50 (w/w) mixture of poly-L-lysine graft copolymers of polyethylene glycol (PLL-PEG) that were modified with an RGD-peptide and a terminal methoxy group, respectively (*33*), to block unspecific binding of labelled nanobodies to the cover slide. Microscopy experiments were performed in presence of an oxygen-scavenging system composed of glucose oxidase (4.5 U*mL^-1^), catalase (540 U*mL^-1^), glucose (4.5 mg*mL^-1^), ascorbic acid (1 mM) and methyl viologen (1 mM) (*65*) to increase photostability. GP130 at the cell surface was labelled by co-incubation with 3 nM of anti-ALFA nanobody site-specifically conjugated with ATTO Rho11 (Rho11) and ATTO 643 (AT643) maleimide (ATTO-TEC), respectively. Alternatively, endogenous GP130 was labelled by co-incubation with 5 nM of HypIL-6 site-specifically conjugated with CoA-DY547-P1 and CoA-DY647-P1 (Dyomics), respectively. Both nanobodies were kept in solution throughout the experiment to ensure consistent high DOL. Cells were imaged without stimulation and after addition of 50 nM HypIL-6. Videos of viable cells showing typical surface densities of 0.1-0.8 copies/µm² per channel were recorded at 30 fps for typically 150 consecutive frames using CellSens 2.2 (Olympus) as acquisition software. Microscopy image stacks were subjected to single-molecule co-localization and co-tracking analysis using a custom-made MATLAB script called SLIMfast 4C as described with more detail in (*42*). Relative co-diffusion levels were determined for each cell recorded in the experiments and corrected for stochastic dual colour labelling. Statistical analysis was performed from typically 15 cells recorded for each condition.

