## Supplementary figures and images for "Differential functional coupling in Gp130-JAK complexes expands the plasticity of the interleukin-6 signaling axis"

### Sup Figures

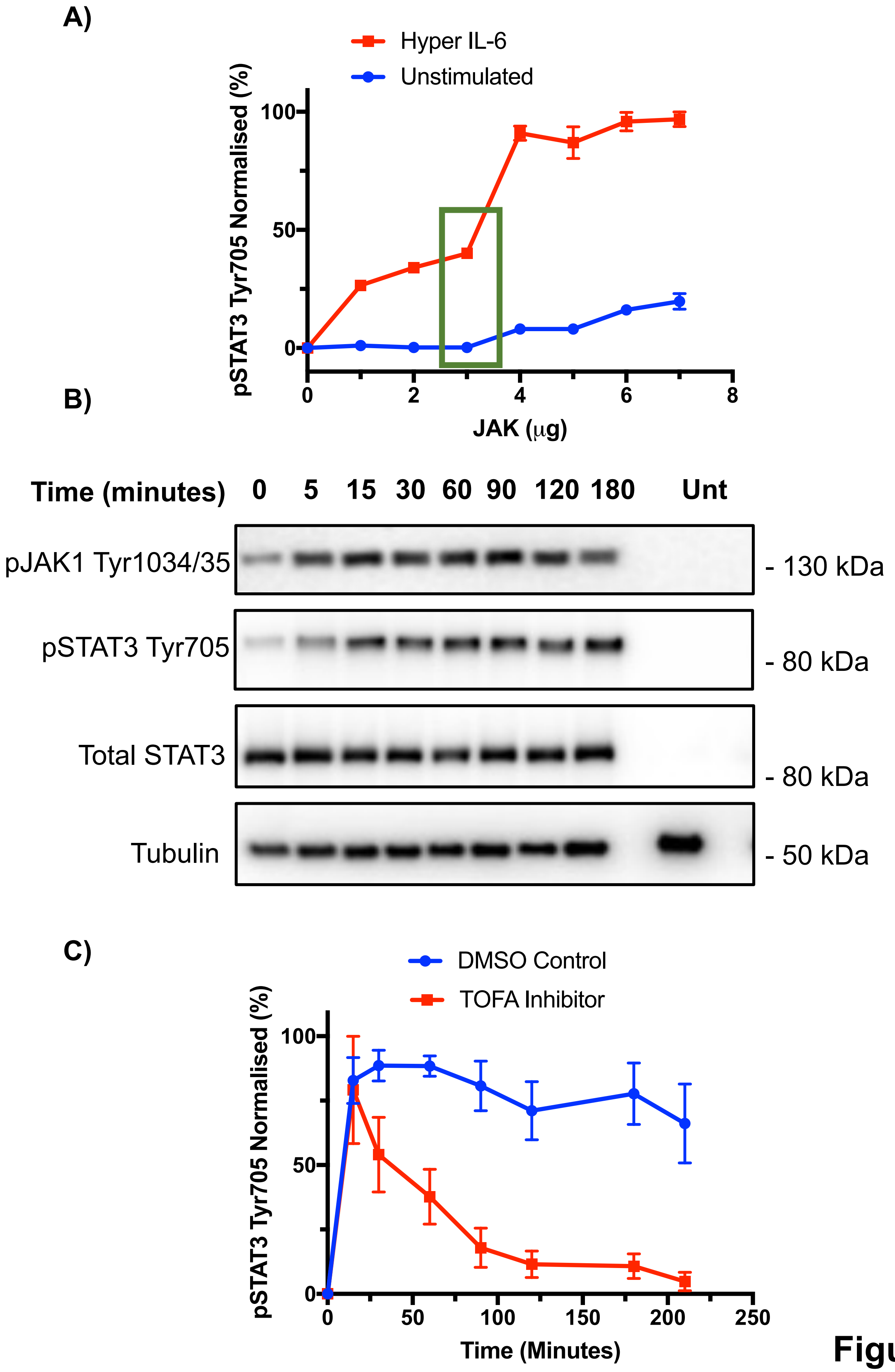

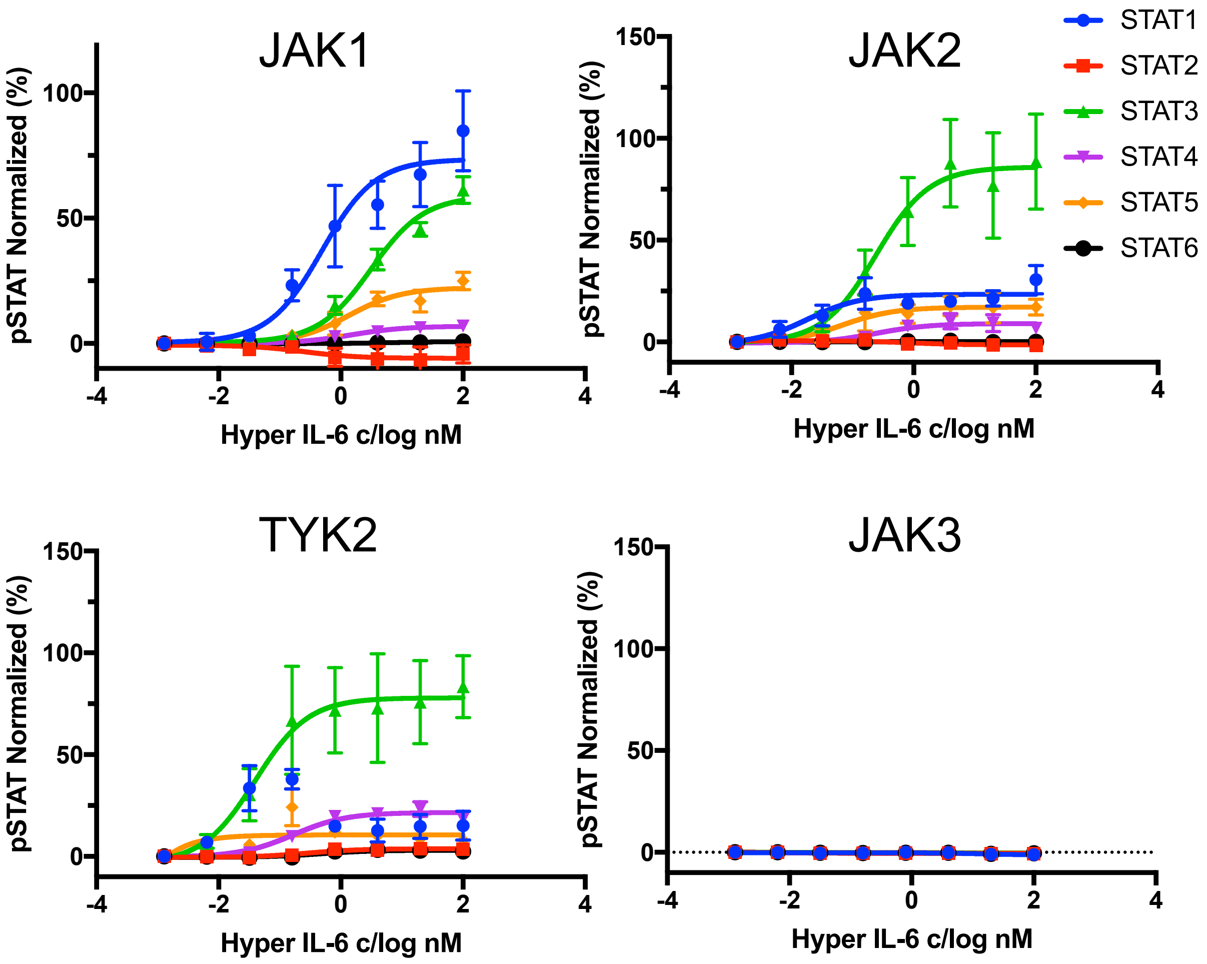

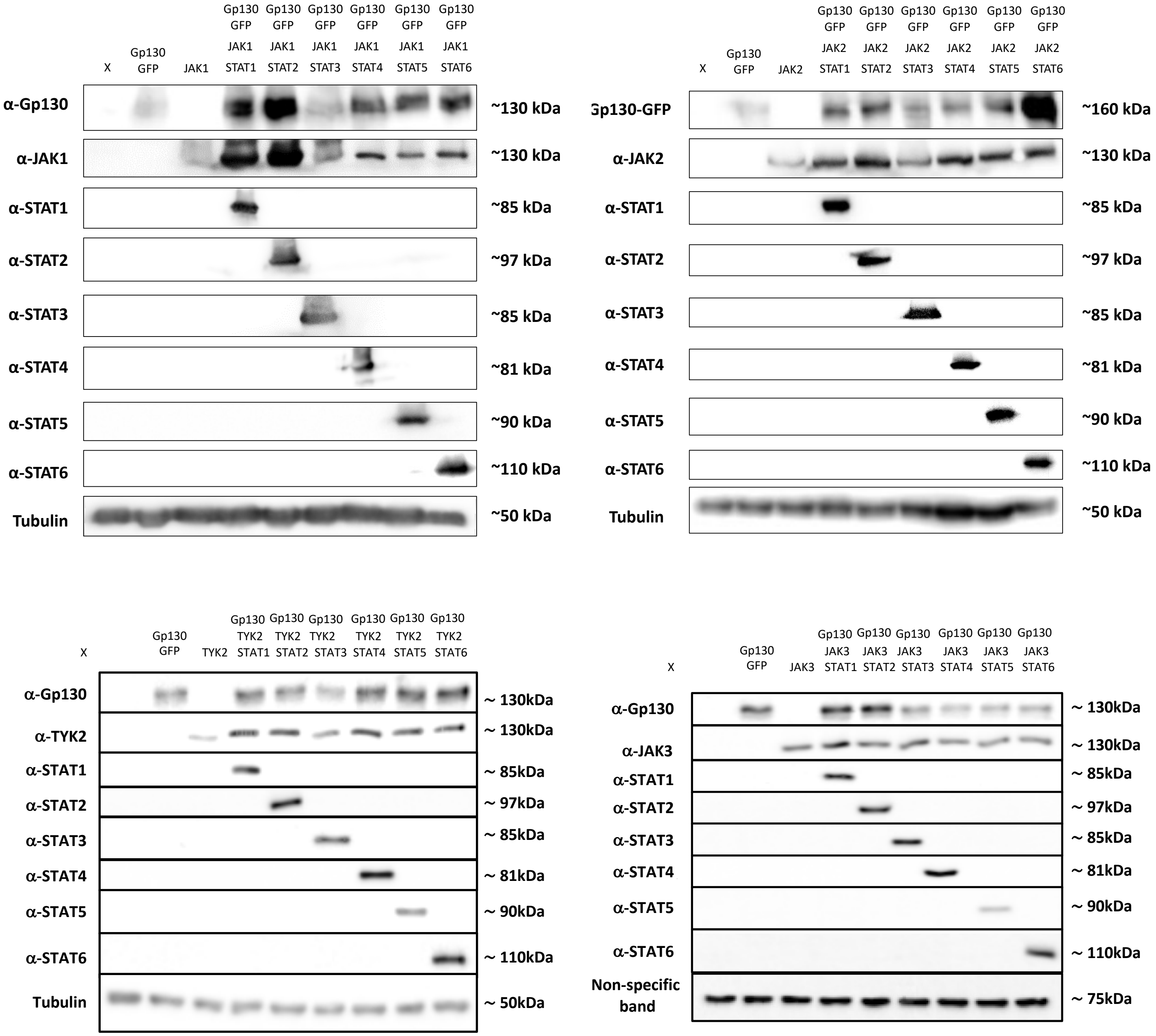

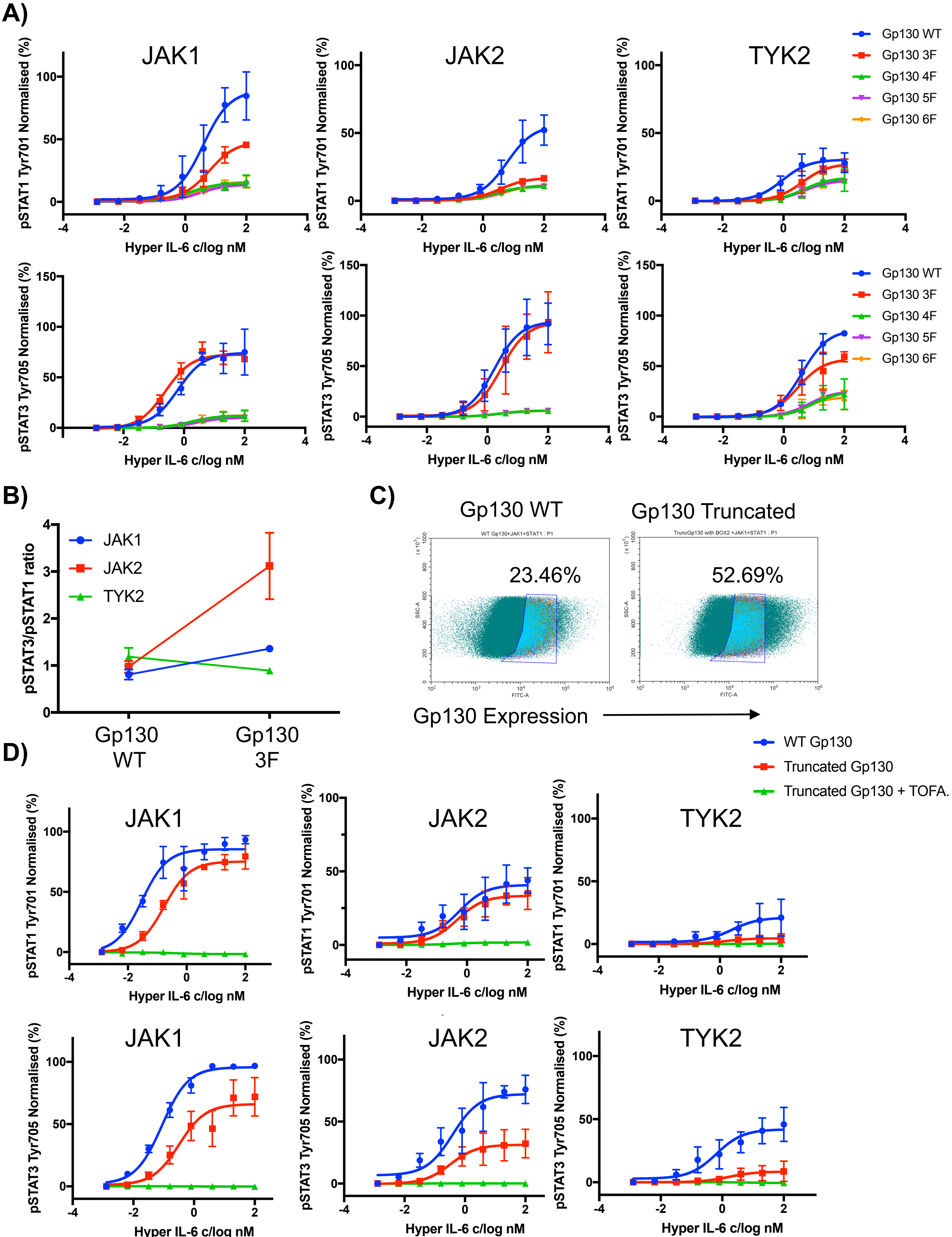

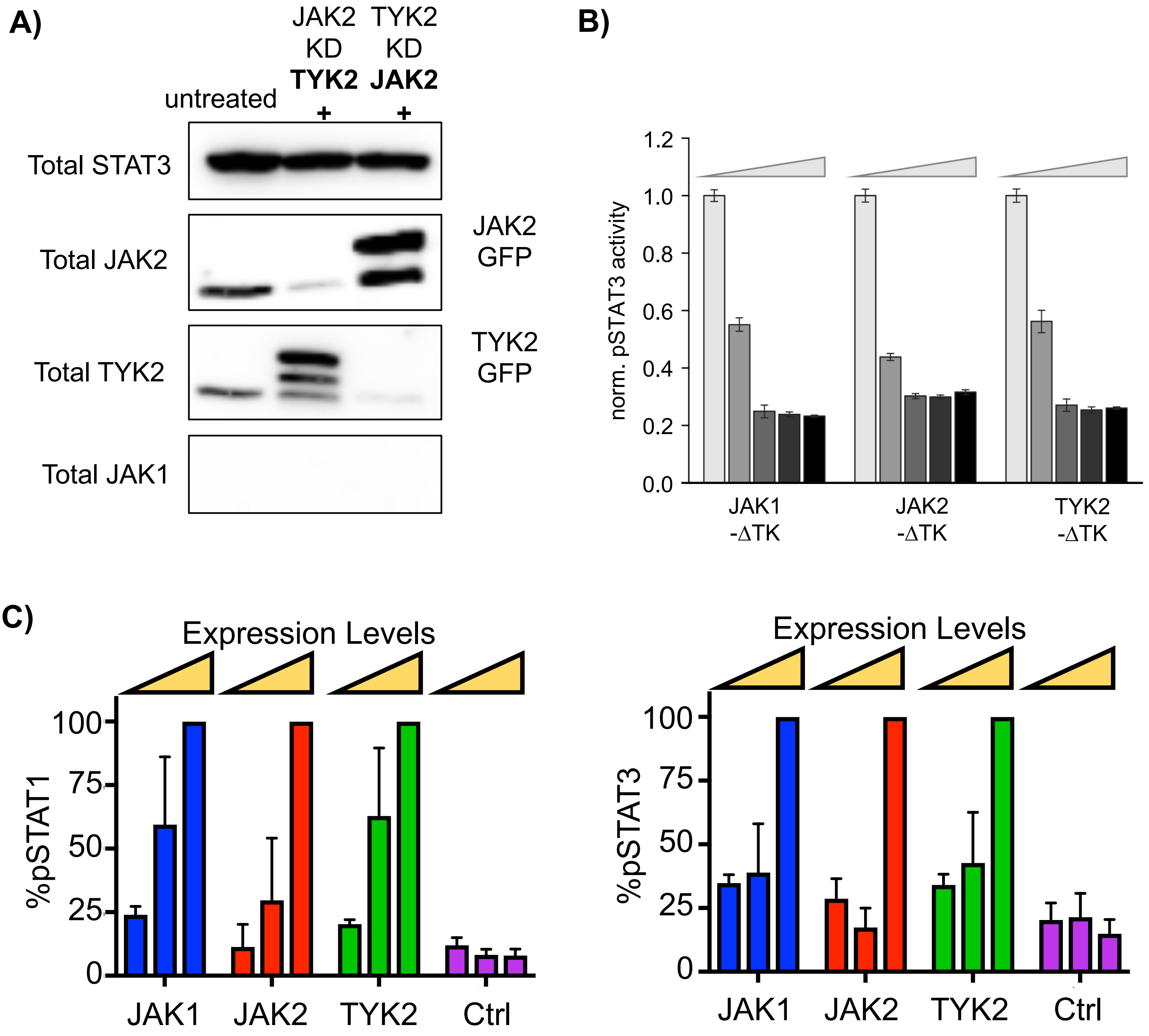

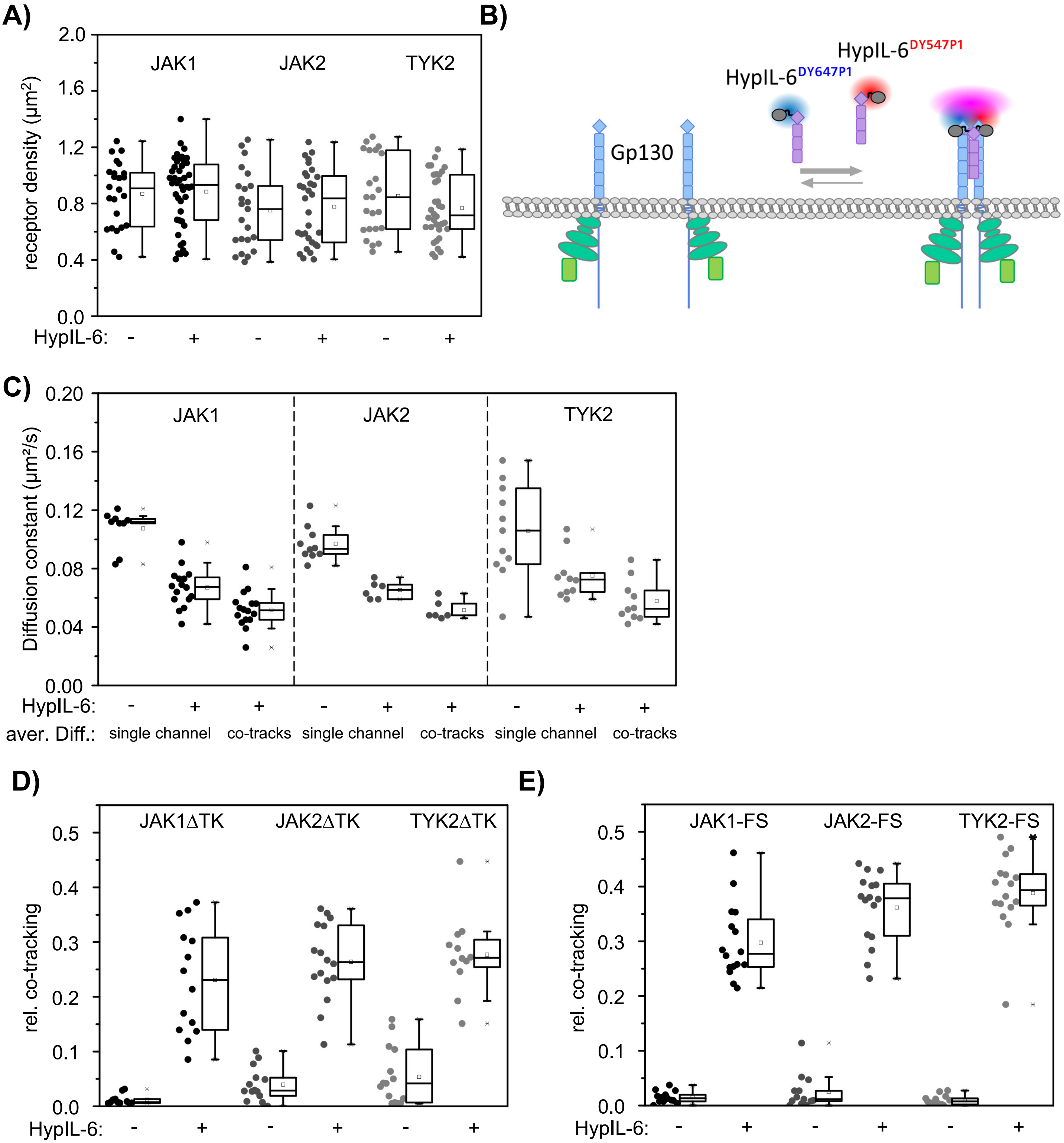

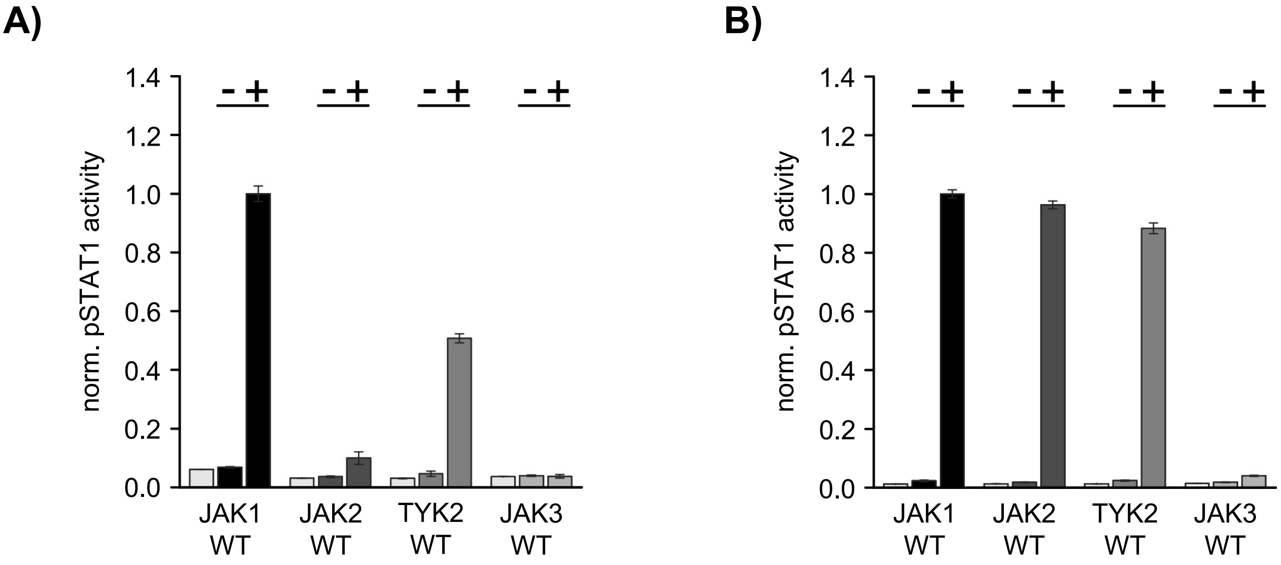
